# Glycosylphosphatidylinositol Biosynthesis and Remodeling are Required for Neural Crest Cell, Cardiac and Neural Development

**DOI:** 10.1101/513507

**Authors:** Marshall Lukacs, Tia Roberts, Praneet Chatuverdi, Rolf W. Stottmann

## Abstract

The glycosylphosphatidylinositol (GPI) anchor attaches nearly 150 proteins to the cell surface. Patients with pathogenic variants in GPI biosynthetic pathway genes display an array of phenotypes including seizures, developmental delay, dysmorphic facial features and cleft palate. There is virtually no mechanism to explain these phenotypes. we identified a novel mouse mutant (*cleft lip/palate, edema and exencephaly; Clpex*) with a hypomorphic mutation in *Post-Glycophosphatidylinositol Attachment to Proteins-2 (Pgap2). Pgap2* is one of the final proteins in the GPI biosynthesis pathway and is required for anchor maturation. We found the *Clpex* mutation results in a global decrease in surface GPI expression. Surprisingly, *Pgap2* showed tissue specific expression with enrichment in the affected tissues of the *Clpex* mutant. We found the phenotype in *Clpex* mutants is due to apoptosis of neural crest cells (NCCs) and the cranial neuroepithelium, as is observed in the GPI anchored *Folate Receptor 1^-/-^* mouse. We showed folinic acid supplementation *in utero* can rescue the cleft lip phenotype in *Clpex*. Finally, we generated a novel mouse model of NCC-specific total GPI deficiency in the *Wnt1-Cre* lineage. These mutants developed median cleft lip and palate demonstrating a cell autonomous role for GPI biosynthesis in NCC development.

## Introduction

Inherited glycophosphatidylinositol deficiency (IGD) disorders are a class of congenital disorders of glycosylation that affect the biosynthesis of the glycosylphosphatidylinositol (GPI) anchor. The clinical spectrum of IGDs is broad and includes epilepsy, developmental delay, structural brain malformations, cleft lip/palate, skeletal hypoplasias, deafness, ophthalmological abnormalities, gastrointestinal defects, genitourinary defects, heart defects, Hirschsprung’s disease, hyperphosphatasia, and nephrogenic defects [1-4]. However, across all GPI deficiency disorders, the most penetrant defects affect the central nervous system and the craniofacial complex [1–4]. Indeed, automated image analysis was able to predict the IGD gene mutated in each patient from facial gestalt [2]. Interestingly, facial gestalt was a better predictor of patient mutation than analysis of the degree of GPI biosynthesis by flow cytometry. Little is known about the mechanism(s) that causes these phenotypes or why disparate tissues are differentially affected[4][4]. We sought to determine the mechanism responsible for these phenotypes using a novel mouse model of reduced enzymatic function within the GPI biosynthesis pathway.

The GPI anchor is a glycolipid added post-translationally to nearly 150 proteins which anchors them to the outer leaflet of the plasma membrane and traffics them to lipid rafts [1]. The biosynthesis and remodeling of the GPI anchor is extensive and requires nearly 30 genes [1]. Once the glycolipid is formed and transferred to the C-terminus of the target protein by a variety of Phosphophatidylinositol Glycan (PIG proteins), it is transferred to the Golgi Apparatus for remodeling by Post-GPI Attachment to Proteins (PGAP proteins). One of these PGAP proteins involved in remodeling the GPI anchor is Post-Glycosylphosphatidylinositol Attachment to Proteins 2 (PGAP2). PGAP2 is a transmembrane protein that catalyzes the addition of stearic acid to the lipid portion of the GPI anchor and cells deficient in *Pgap2* lack stable surface expression of a variety of GPI-anchored proteins (GPI-APs) [1,5]. Autosomal recessive mutations in *PGAP2* cause Hyperphosphatasia with Mental Retardation 3 (HPMRS3 OMIM # 614207), an IGD that presents with variably penetrant hyperphosphatasia, developmental delay, seizures, microcephaly, heart defects, and a variety of neurocristopathies including Hirschsprung’s disease, cleft lip, cleft palate, and facial dysmorphia [5–8]. Currently, there is no known molecular mechanism to explain the cause of these phenotypes or therapies for these patients.

In a forward genetic ENU mutagenesis screen, we previously identified the *Clpex* mouse mutant with Cleft Lip, Cleft Palate, Edema, and Exencephaly *(Clpex)* [9]. Here we present evidence that this mutant phenotype is caused by a hypo-morphic allele of *Pgap2*. To date, embryonic phenotypes of GPI biosynthesis mutants have been difficult to study due to the early lethal phenotypes associated with germline knockout of GPI biosynthesis genes [10–13]. The International Mouse Phenotyping Consortium labelled *Pgap2* homozygote knockout mice preweaning lethal and the Deciphering the Mechanisms of Developmental Disorders initiative labelled *Pgap2* homozygotes early lethal as no embryos were recovered at embryonic day E9.5. In this study, we took advantage of the *Clpex* hypomorphic mutant to determine the mechanism of the various phenotypes and tested the hypothesis that GPI-anchored Folate Receptor 1 (FOLR1) is responsible for the phenotypes observed.

As we observed tissue specific defects in NCCs in the *Clpex* germline mutant, we sought to determine the cell autonomous requirement for GPI biosynthesis pathway generally in NCC development. To do this, we took a conditional approach to abolish GPI biosynthesis in NCCs using the *Wnt1*-Cre transgene and a floxed allele of a critical initiator of GPI biosynthesis. The *Clpex* mutant and our NCC conditional mutant serve as models for studying the effect of GPI biosynthesis defects in development of the craniofacial complex and testing potential therapeutics *in utero* for patients.

## Results

### The *Clpex* mutant phenotype is caused by a missense mutation in *Pgap2*

We previously identified the *Clpex (cleft lip and palate, edema, and exencephaly)* mutant in a mouse N-ethyl-N-nitrosourea (ENU) mutagenesis screen for recessive alleles leading to organogenesis phenotypes [9]. *Clpex* homozygous mutants displayed multiple partially penetrant phenotypes. In a subset of 70 mutants from late organogenesis stages (~E16.5-E18.5), we noted cranial neural tube defects (exencephaly) in 61 (87%), cleft lip in 22 (31%), cleft palate in 13 (19%), and edema in 6 embryos (9%) (Fig. 1A-H). Skeletal preparations of *Clpex* mutants identified a defect in frontal bone ossification (Fig. 1I-L, n=5/5 mutants) and a statistically significant decrease in limb length (Fig. 1M-P). We previously reported a genetic mapping strategy with the Mouse Universal Genotyping Array which identified a 44 Mb region of homozygosity for the mutagenized A/J genome on chromosome 7 (Fig. 1Q) [9]. We then took a whole exome sequencing approach and sequenced 3 *Clpex* homozygous mutants. Analysis of single base pair variants which were homozygous in all 3 mutants with predicted high impact and not already known strain polymorphisms in dbSNP left only one candidate variant (Table 1). This was a homozygous missense mutation in the initiating methionine (c.A1G, p.M1V) in exon 3 of *post-GPI attachment to proteins 2 (Pgap2)*. We confirmed the whole exome sequencing result by Sanger Sequencing (Fig. 1R). This mutation abolishes the canonical translation start codon for *Pgap2*. However, there are alternative start sites of *Pgap2* and multiple alternatively spliced transcripts that may lead to production of variant forms of *Pgap2*.

**Figure 1.**
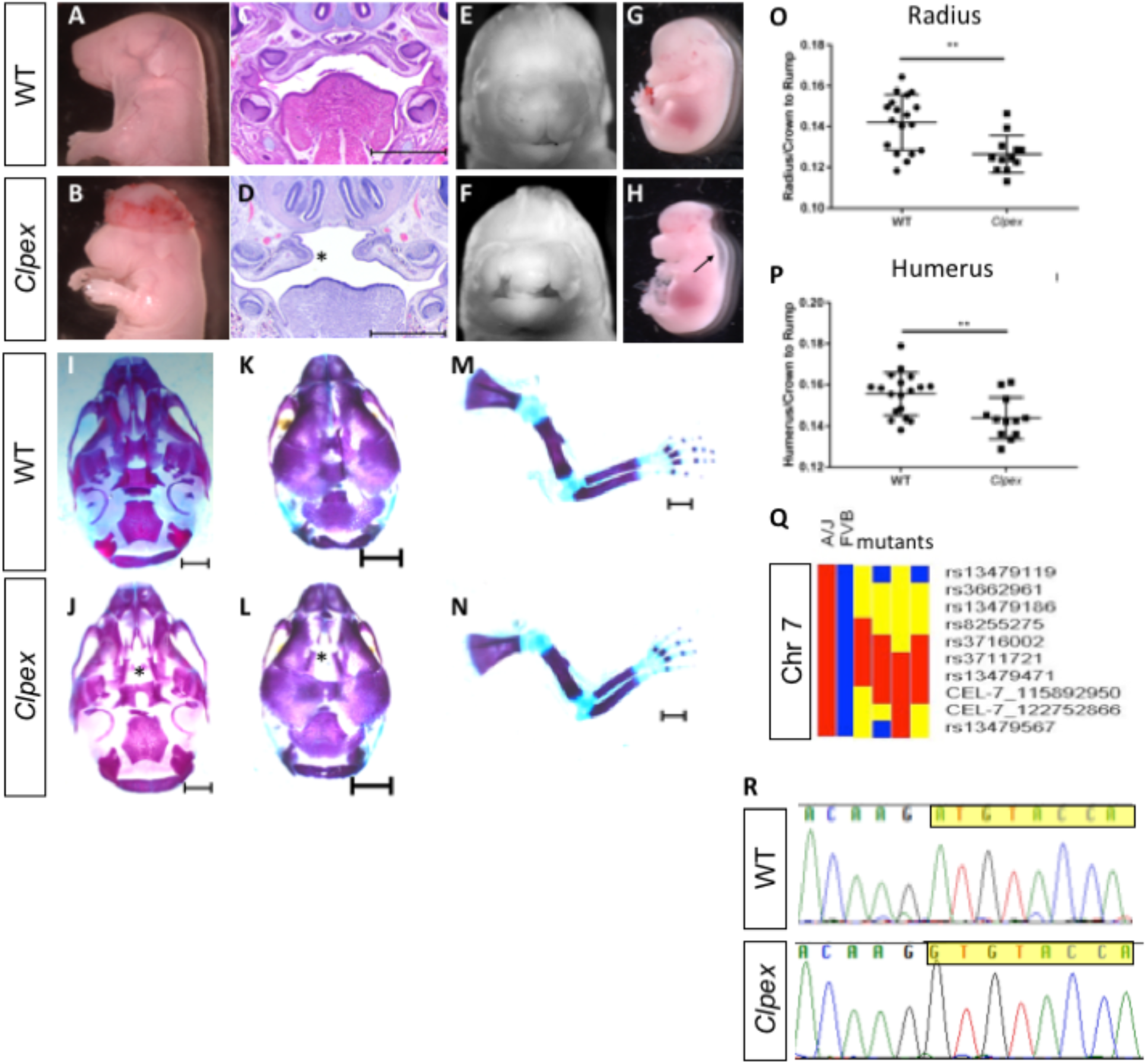
The *Clpex* mutant phenotype is caused by a hypomorphic mutation in *Pgap2*. Whole mount E18.5 (A,E) and E15.5 (G) WT embryos. Whole mount E18.5 (B, F) and E15.5 (H) *Clpex* mutant embryos. H&E staining of WT E15.5 (C) and *Clpex* (D) coronal sections. Skeletal preparation of WT skull ventral view (I), dorsal view (K). Skeletal preparation of *Clpex* mutant skull ventral view (J) and dorsal view (L). Asterix indicates absent palatine bone in mutant. Skeletal preparation of WT limb (M), and *Clpex* mutant limb (N). Quantification of WT and mutant radial (O) and humeral (P) length normalized to the crown to rump ratio. Mapping data for *Clpex* mutation (Q). Sanger sequencing of *Pgap2* exon 3 in WT and *Clpex* mutant with exon 3 highlighted starting at the initiating methionine.(R). Scale bar indicates 500μm in C,D and 1mm in I-N. (** p< 0.001).

**Table 1.** Exome analysis identifies variant in *Pgap2*.

| Variant Filters | Number |
| --- | --- |
| Total variants | 145,956 |
| Homozygous in all 3 mutants | 120,393 |
| Chromosome 7 | 5,854 |
| 81-125 Mb | 2,196 |
| “High” impact | 9 |
| Not in dbSNP | 7 |
| Single base pair change | 1: *Pgap2* |

To determine whether the *Clpex* phenotype was caused by the missense mutation in *Pgap2*, we performed a genetic complementation test using the *pgap2^tm1a(EUCOMM)Wtsi^* (hereafter referred to as *Pgap2^nul1^)* conditional gene trap allele. We crossed *Pgap2^Clpex/+^* heterozygotes with *Pgap2^null/+^* heterozygotes to generate *pgap2^Clpex/nul1^* embryos. *Pgap2^Clpex/nul1^* embryos displayed neural tube defects, bilateral cleft lip, and edema similar to the *pgap2^Clpex/Clpex^* embryos at E13.5 (Fig. 2 A-H). *pgap2^Clpex/nul1^* embryos also displayed micro-opthalmia (Fig. 2F) and more penetrant cleft lip and edema phenotypes than observed in *pgap2^Clpex/Clpex^* homozygotes (Fig. 2I). *Pgap2^Clpex/nul1^* embryo viability was decreased in this line with lethality at approximately E13.5-E14.5, precluding analysis of palatal development in these mutants. Histological analysis of the heart in *pgap2^Clpex/nul1^* E13.5 embryos showed pericardial effusion, a reduction in thickness of the myocardium, and an underdeveloped ventricular septum and valves (Fig. 2J-Q). As the *Clpex* allele failed to complement a null allele of *Pgap2*, we concluded the *Clpex* phenotype is caused by a hypomorphic allele of *Pgap2*. The Deciphering Mechanisms of Developmental Disorders (DMDD) initiative has generated *Pgap2^null/null^* homozygotes and classified them as “early lethal” abnormal chorion and trophoblast layer morphology [12]. These data argue *Pgap2* is required for early embryogenesis and our *Clpex* allele must be hypomorphic, as the embryos survive to E18.5.

**Figure 2.**
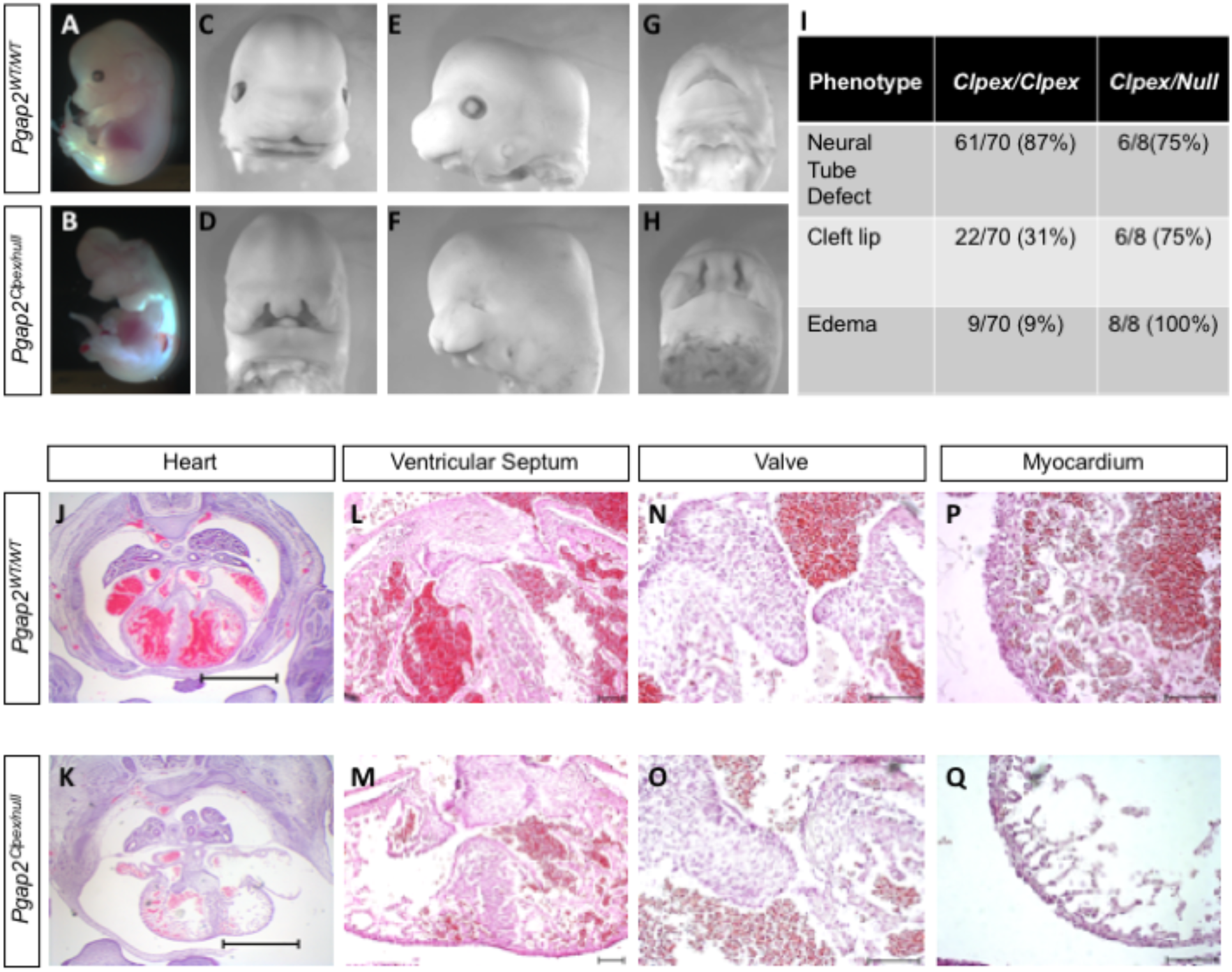
*Pgap2^nul1^* allele fails to complement *Pgap2^Clpex^* allele. Whole mount image of E13.5 WT (A,C,E,G) and *Pgap2^Clpex/null^* mutant (B,D,F,H). Penetrance of some key phenotypes is compared in I. Cardiac histology of E14.5 WT (J) and *Pgap2^Clpex/null^* mutant (K,), scale bar indicates 1 mm. Higher power images of the ventricular septum (L, M), valve (N, O), and myocardial wall (P, Q). Scale bar indicates 1mm in J,K and 100μM in L-Q.

### *Pgap2* is dynamically expressed throughout development

Based on the tissues affected in the *Clpex* mutant, we hypothesized *Pgap2* is expressed in the neural folds and facial primordia of the developing mouse embryo at early stages. We used the lacZ expression cassette within the *Pgap2^null^* allele to perform detailed expression analysis of *Pgap2* throughout development (n= 21 litters at multiple developmental stages). We performed RNA *in situ* hybridization in parallel for some stages to test the fidelity of the lacZ expression and found high concordance. *Pgap2* was expressed relatively uniformly and ubiquitously at neurulation stages in the mouse from E7.5-E8.5 (Fig. 3A-D). We also noted extraembryonic expression at E7.5, consistent with the abnormal placental development observed in *Pgap2^null/null^* embryos (Fig. 3A,B) [12]. At E9.5-11.5, there was clear enrichment of *Pgap2* expression in the first branchial arch (Fig. 3E-G). *Pgap2 in situ* hybridization identified a similar pattern of expression as observed in the *Pgap2* LacZ reporter allele at E9.5 (Fig.3E) At E10.5 and E11.5, expression was enriched in the limb bud, somites, first branchial arch, eye, forebrain and midbrain (Fig. 3F-J). There was also increased expression at the medial aspects of both medial and lateral nasal processes at E10.5 (Fig. 3G-H), and strong expression in the heart starting at E11.5 (Fig. 3I,J). Later in organogenesis stages, *Pgap2* showed more regionalized and enriched expression, including in the ganglion cell layer of the retina at E12.5 and E14.5 (Fig. 3K,L). At E16.5 *Pgap2* was expressed in the salivary gland, epidermis, stomach, nasal conchae, myocardium, bronchi, kidney, uroepithelium, lung parenchyma, a specific layer of the cortex, and ear (Fig. 3M-T). Interestingly, *Pgap2* showed lower expression in the liver (Fig. 3S) and most of the brain except for a thin layer of the cortex and the choroid plexus at E16.5 (Fig. 3S, U-V, X). We also noted expression in the genital tubercle (Fig. 3W). We conclude *Pgap2* shows tissue specific regions of increased expression which may help to explain why certain tissues such as the craniofacial complex, central nervous system, and heart are differentially affected in GPI biosynthesis mutants. These data are in contrast to previous reports where some GPI biosynthesis genes are shown to be ubiquitously and uniformly expressed, including *Pign* in the mouse and *pigu* in zebrafish [10, 14]. Our *Pgap2* expression is more consistent with the expression of *Pigv* which is enriched in *C. elegans* epidermal tissues [15]

**Figure 3.**
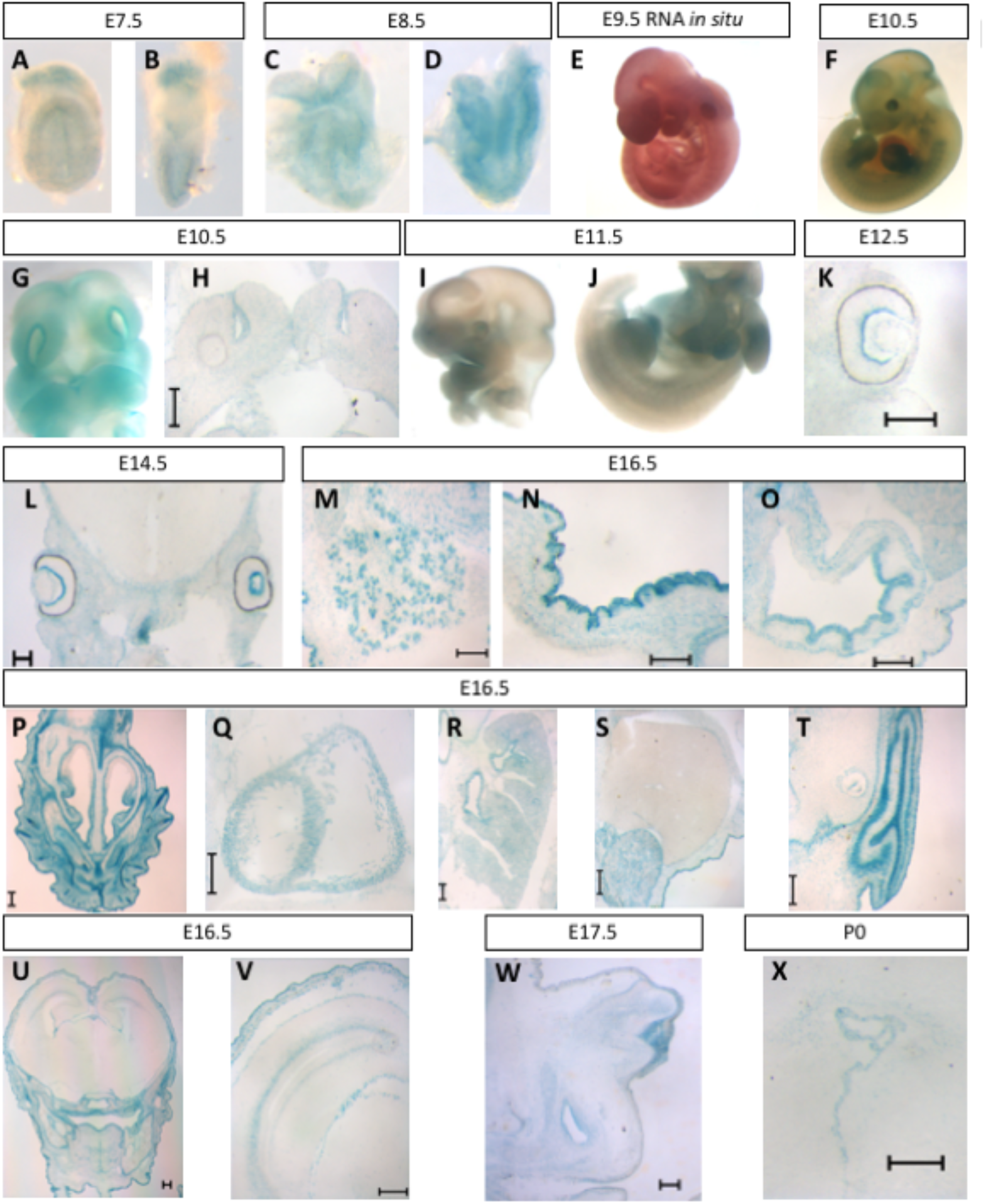
*Pgap2* is dynamically expressed throughout embryogenesis. Whole mount *Pgap2* Xgal staining in E7.5 (A-B), E8.5 (C-D), E10.5 (F-G) E11.5 (I-J). *Pgap2 in situ* hybridization at E9.5 (E). Transverse section through the anterior facial tissues at E10.5 (H) at the future site of lip closure. (K-X) Xgal section staining is shown from embryos at E12.5, E14.5, E16,5 E17.5 and P0. Expression is seen in the ganglion cell layer of the retina (K,L), salivary gland (M), epidermis (N), stomach (O), nasal conchae (P), myocardium (Q), lung parenchyma (R), kidney (S), ear (T), cerebral cortex (U-V), genital tubercle (W) and brain/choroid plexus (X). Scale bar indicates 200 μm.

### *Pgap2* is required for the proper anchoring of GPI-APs, including FOLR1

PGAP2 is the final protein in the GPI biosynthesis pathway and catalyzes the addition of stearic acid to the GPI anchor [16]. GPI is a glycolipid added post-translationally to nearly 150 proteins in the endoplasmic reticulum (ER) and remodeled in the Golgi Apparatus. GPI anchored proteins (GPI-APs) require the GPI anchor for their presentation in the outer leaflet of the plasma membrane and association with lipid rafts [1, 16]. In the absence of *Pgap2*, cells lack a variety of GPI-APs on the cell surface leading to a functional GPI deficiency [5–8, 16]. To determine the effect of the *Clpex* mutation on *Pgap2* function, we performed Fluorescein-labeled proaerolysin (FLAER) flow cytometry staining to quantify the overall amount of the GPI anchor on the cell surface. FLAER is a bacterial toxin conjugated to fluorescein that binds directly to the GPI anchor in the plasma membrane. We hypothesized *Pgap2* function is impaired in *Clpex* mutants due to the ENU mutation in the initiating methionine. We found mouse embryonic fibroblasts (MEFs) from *Clpex* mutants displayed a significantly decreased expression of FLAER compared to wildtype MEFs, consistent with a defect in GPI biosynthesis (n=3 separate experiments with 4 WT and 4 *Clpex* cell lines) (Fig. 4A,B).

**Figure 4.**
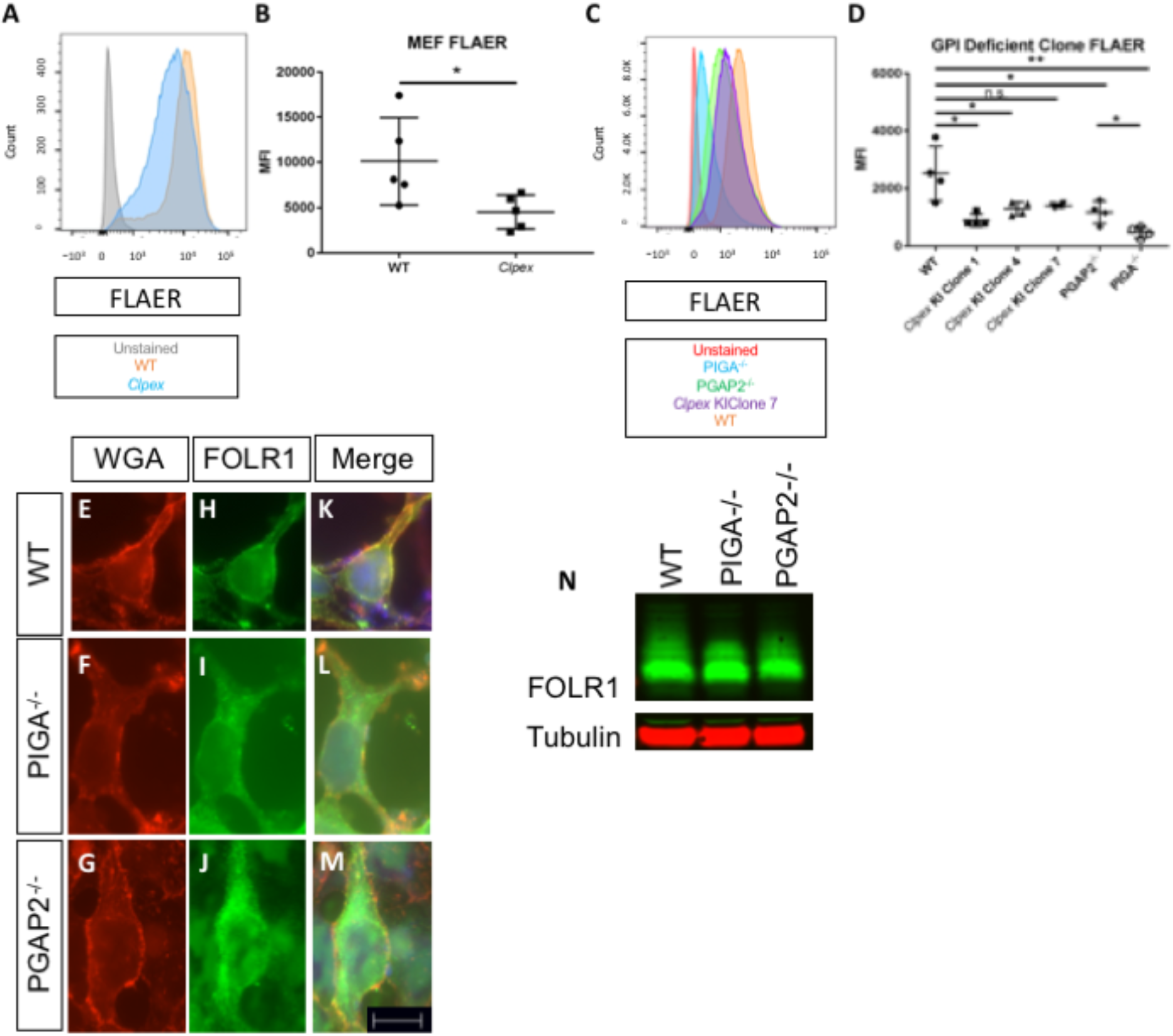
*Pgap2* is required for proper anchoring of GPI-APs including FOLR1. (A) FLAER staining of WT (orange) and *Clpex* (blue) MEFs, (unstained control in grey) with quantification (B). FLAER staining of WT (orange) *Clpex* KI Clone 7 (purple), *PGAP2r^-/-^* (green), *PIGA^-/-^* (blue) HEK293T cells, and unstained control (red) (C) with quantification (D). Wheat germ agglutinin (WGA) staining in WT (E), *PIGA^-/-^* (F), *PGAP2^-/-^* (G) HEK293T cells. FOLR1 myc staining in WT (H), *PIGA^-/-^* (I), *PGAP2^-/-^* (J) HEK293T cells. Merge of WGA and FOLR1 for WT (K), *PIGA^-/-^* (L), and *PGAP2^-/-^* (M). Western blot for αmyc-FOLR1 (green) and αTubulin (red) loading control from cell lysates of WT, *PIGA^-/-^*, and *PGAP2^-/-^* cells. (*p<0.05, **p<0.001, Scale bar indicates 10μM.)

Our genetic complementation analysis results suggested the *Clpex* allele might be a hypomorphic allele of *Pgap2*. To test this hypothesis, we generated human embryonic kidney (HEK) 293T clones with a 121bp deletion in exon 3 of *PGAP2* with CRISPR/Cas9 (termed *PGAP2^-/-^* cells; Fig. S1). In parallel, we recapitulated the *Clpex* mutation in 3 independent clones of HEK293T cells by CRISPR/Cas9 mediated homologous directed repair (termed *Clpex KI* Clones 1, 4, and 7; Fig.S1). We found there was a statistically significant decrease in FLAER staining between WT and 2 of 3 KI clones and the *PGAP2^-/-^* cells. However, we observed no statistical difference in FLAER staining in *PGAP2^-/-^* cells when compared to *Clpex KI* cells (Fig. 4C,D). Therefore, we conclude the *Clpex* missense mutation severely affects PGAP2 function similar to the effect seen upon total depletion of *PGAP2*. As a positive control, we used CRISPR/Cas9 to delete *phosphatidylinositol glycan anchor biosynthesis, class A (Piga;* Fig.S1). *Piga* is the first gene in the GPI biosynthesis pathway and is absolutely required for GPI biosynthesis [6, 17]. We utilized CRISPR/Cas9 to generate a 29bp out-of-frame deletion in exon 3 of *PIGA*. These *PIGA^-/-^* cells showed an even further decrease in FLAER staining compared to *PGAP2^-/-^* cells, confirming our staining accurately reflects GPI anchor levels (n=4 separate experiments; Fig. 3C-D).

Current estimates suggest nearly 150 genes encode proteins which are GPI anchored [18]. Our manual review of the MGI database found 102 GPI-APs have been genetically manipulated and phenotyped in mice [19]. Of these, the null allele of *Folr1* has a phenotype most similar to the *Clpex* mutant with cranial neural tube defects, cleft lip/palate, and heart outflow tract phenotypes [20]. Tashima et. al. previously showed *PGAP2* is required for stable cell surface expression of FOLR1 in CHO cells [16]. To confirm this finding, we overexpressed a myc-tagged FOLR1 construct in WT and *PGAP2r^-/-^* 293T cells and assessed the presentation on the plasma membrane by immunocytochemistry for wheat germ agglutinin (WGA). We observed a decrease in co-localization of FOLR1 with WGA in *PGAP2r^-/-^* cells compared to controls (Fig. 4E,F,H,I,K,L). In the absence of *PIGA*, cells lack the surface expression of any GPI-APs [1]. We found *PIGA^-/-^* cells showed decreased co-localization of FOLR1 with WGA similar to *PGAP2^-/-^* cells (representative images from n=5 separate experiments) (Fig. 4G,J,M). However, both *PIGA^-/-^* and *PGAP2^-/-^* cells produced similar amounts of FOLR1 protein by western blot indicating that the defect is in trafficking, and not protein production (Fig. 4N).

### Neural crest cells and cranial neuroepithelium display increased apoptosis in the *Clpex* mutant

A number of GPI-APs are critical for cranial neural crest cell (cNCC) migration and survival; including ephrins and FOLR1 [21–29]. This led us to the hypothesis that cNCC migration may be impaired in *Clpex* mutants ultimately causing the cleft lip and palate phenotype [22]. To test whether cNCC migration was impaired in the *Clpex* mutant, we performed a NCC lineage trace using the *Wnt1-Cre* (*B6.Cg-H2afv^Tg(Wnt1-cre)11Rth^Tg(Wnt1-* GAL4)11Rth/J) in combination with the R26R LacZ reporter (B6.129S4-*Gt(ROSA)26Sor^tm1Sor^/J* (R26R^Tg^) to create *Wnt1-Cre;R26R^Tg/wt^;Pgap2^clpex/clpex^* mutants in which the NCCS are indelibly labeled with LacZ expression at E9.5 and E11.5 [30–34]. We observed no significant deficit in cNCC migration in the mutant embryos as compared to littermate controls at either stage (representative images of n=2 E9.5 mutants and n=5 E11.5 mutants) (Fig. 5A-F). However, we observed hypoplasia of the medial and lateral nasal processes at E11.5, suggesting the *Clpex* phenotype is due to earlier defects in NCC survival (Fig 5E-F). As *Pgap2* was highly expressed in the epithelium and epithelial barrier defects are a known cause of cleft palate, we next sought to determine whether the epidermis was compromised in the Clpex mutant [35]. We performed a Toluidine Blue exclusion assay but found no significant defects in barrier formation in the mutant (Fig. S2).

**Figure 5.**
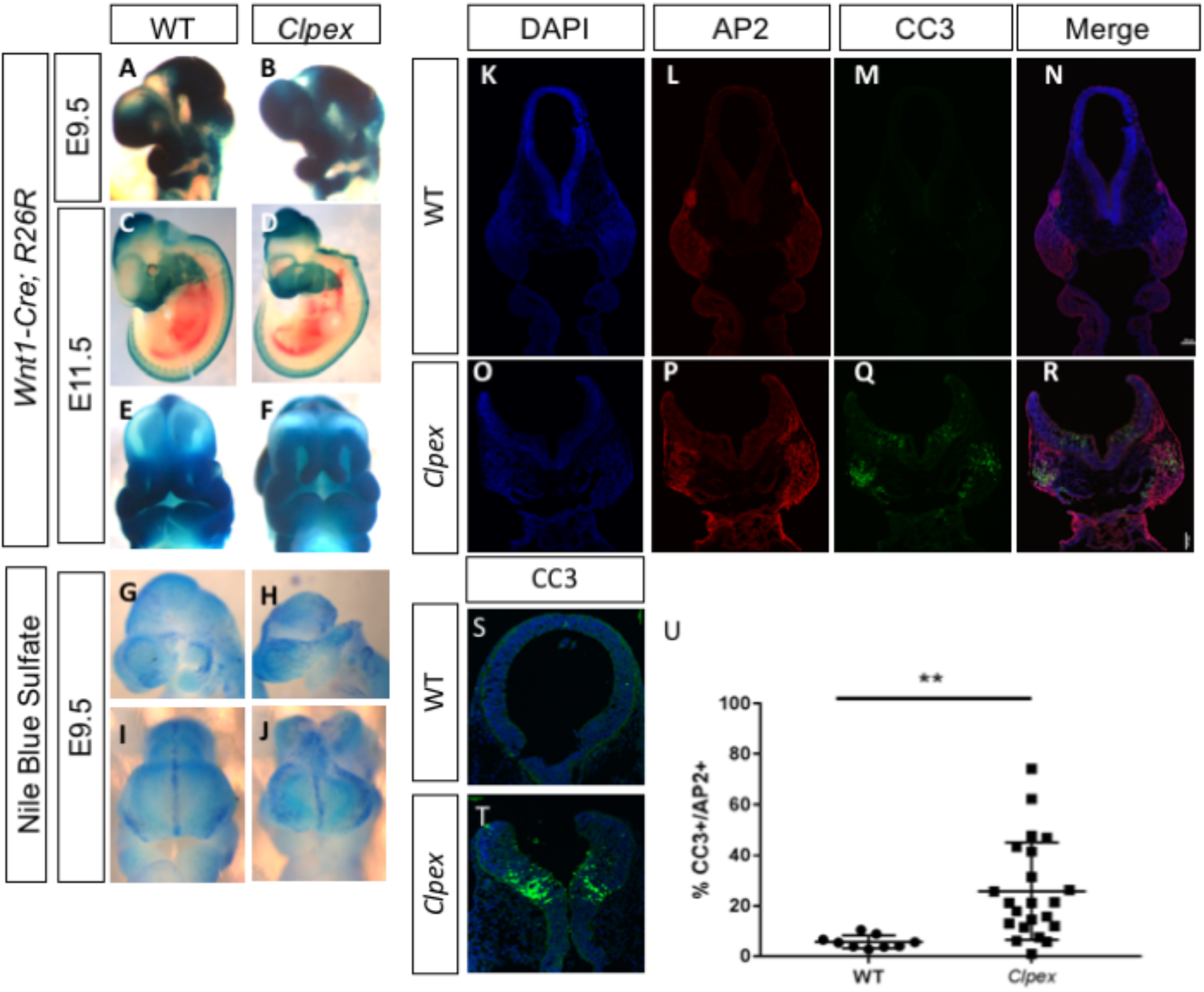
*Clpex* cNCCs and neuroepithelium undergo apoptosis at E9.5. *Wnt1-Cre*, R26R NCC lineage trace in WT (A, C, E) and *Clpex* mutant (B,D,F) at E9.5 (A,B) and E11.5 (C-F). Nile Blue Sulfate whole mount viability stain in WT (G, I) and *Clpex* mutant (H,J) E9.5 embryos. WT E9.5 embryo stained for DAPI (K), AP2 (L) CC3 (M; merged image in N). *Clpex* E9.5 embryo stained for DAPI (O), AP2 (P) CC3 (Q; merged image in R). Higher power image of WT (S) and *Clpex* mutant (T) neuroepithelium stained with CC3 and DAPI. Quantification of CC3+ cells over AP2+ cells in the first branchial arch Region of Interest (U). **p<0.001. Scale bar indicates 100μm.

*Folr1^-/-^* mice and zebrafish *Folr1* morphants display increased cell death and decreased proliferation in the facial primordia [26, 28, 29, 36]. We hypothesized a similar mechanism may be responsible for the cleft lip and palate in the *Clpex* mutant embryos. To test this hypothesis, we performed whole mount Nile Blue Sulfate viability staining in WT and homozygous *Clpex* mutant embryos at E9.5, a stage before we observed hypoplasia of the nasal processes. We found there was an increase in cell death in the facial primordia, including the frontonasal prominence, the first brachial arch, and cranial neuroepithelium in *Clpex* mutants as compared to controls (representative images from n=12 mutants, 5 separate experiments) (Fig. 5G-J). To confirm this result and determine which cell population was undergoing apoptosis in the *Clpex* mutants, we performed immunohistochemistry for αAP2 to mark NCCs and the apoptosis marker Cleaved Caspase 3 (CC3). We found the cNCCs of the first arch and a specific population of cells within the neuroepithelium were undergoing apoptosis significantly more frequently in *Clpex* homozygous mutants (Fig. 5K-R). The ratio of CC3-positive to AP2-positive cells revealed a highly significant increase in the percentage of CC3-positive cells in the first arch of *Clpex* mutants (n= 2 or more sections from 3 WT and 6 *Clpex* mutants) (Fig. 5U) We also observed apoptosis in the cranial neuroepithelium at the dorsolateral hinge points (Fig. 5S-T). The dorsolateral hingepoints constrict bilaterally in order to close the neural tube. This apoptosis was exclusively confined to the cranial aspects of the neural tube at the midbrain-hindbrain boundary

### Dietary folinic acid supplementation partially rescues cleft lip in *Clpex* mutants

Dietary folinic acid supplementation has been shown to rescue the early embryonic lethal phenotype of *Folr1^-/-^* mice and these mice can then survive to adulthood [26, 28, 37]. Our data suggest FOLR1 receptor trafficking is impaired in the *Clpex* mutant (Fig. 4), leading us to hypothesize folinic acid supplementation *in utero* may rescue the *Clpex* phenotype. We further hypothesized the folinic acid diet would have a greater beneficial effect as folinic acid (reduced folate) has a higher affinity for other folate receptors including *solute carrier family 19 (folate transporter), member 1 (Slc19a1)* and *solute carrier family 46, member1 (Slc46a1),)* which are not GPI anchored [38]. In comparison, folic acid has a higher affinity for the GPI anchored folate receptors FOLR1 and FOLR2 [39, 40]. We supplemented pregnant *Clpex* dams from E0-E9.5 or E16.5 with a 25 parts per million (ppm) folinic acid diet, 25 ppm folic acid diet or control diet, and collected *Clpex* mutants for phenotypic analysis and Nile Blue staining for cell viability. We first found folinic acid supplementation decreased the Nile Blue staining for non-viable cells in the facial primordia of E9.5 mutants compared to controls (n=2 mutants; Fig. 6A,B). We also found the folinic acid diet had an effect on *Clpex* phenotypes at E16.5. The treatment group had a significantly smaller proportion of mutants with cleft lip (p=0.02) but there was no effect on the incidence of NTD or cleft palate (Fig. 6C,D). We did note a decrease in mutants with edema, however, this decrease was not statistically significant given our sample size (p=0.06; Fig. 6C,D). Consistent with our hypothesis, we found the folinic acid reduced the number of mutants with cleft lip by 23% (2/25 mutant vs. 22/70 control), which is more significant than the 10% reduction observed in folic acid treated mice (3/14, Fig. 6C,D). Therefore, we conclude folinic acid treatment increased the viability of *Clpex* facial primordia and decreased the incidence of cleft lip among *Clpex* mutants.

**Figure 6.**
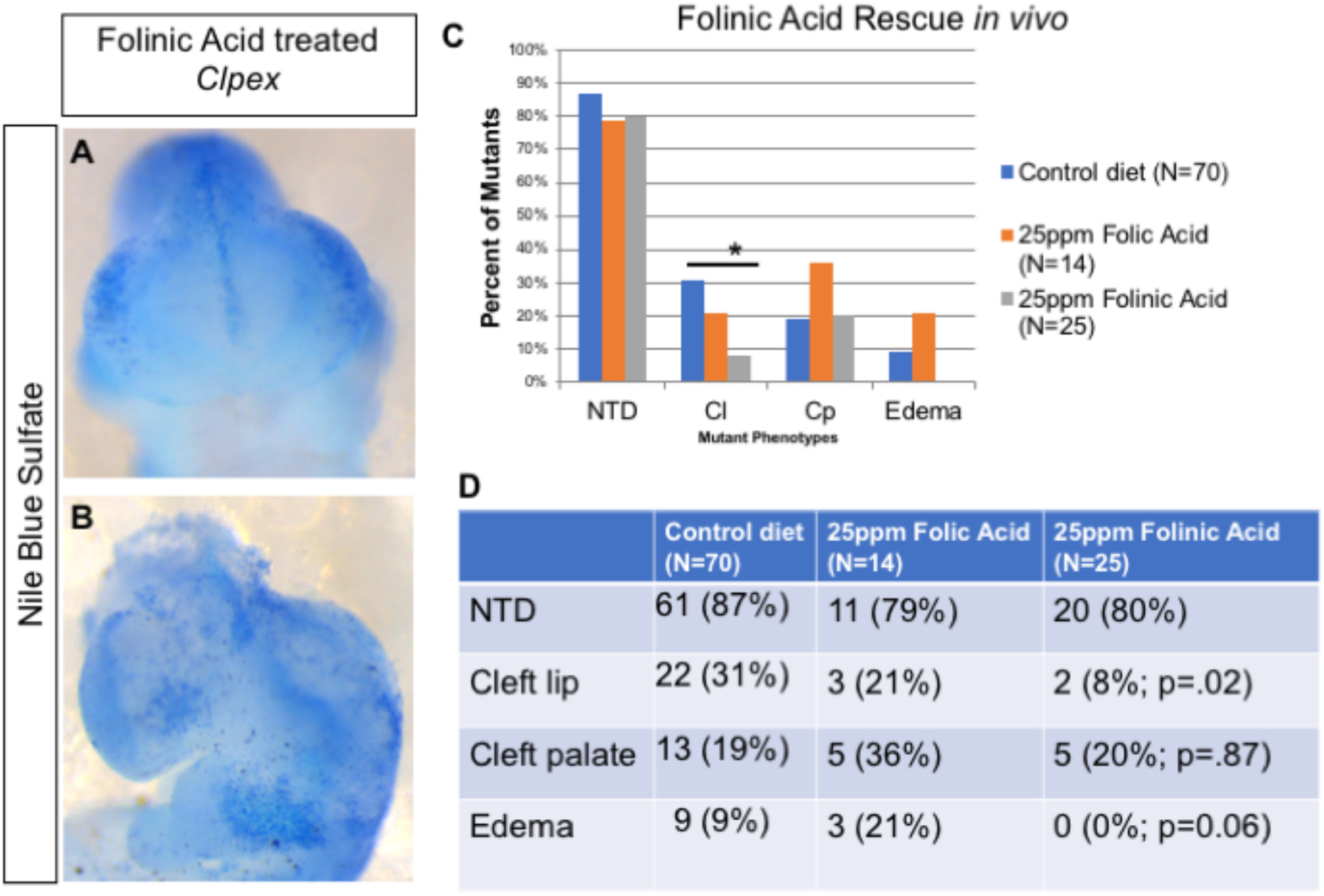
Folinic Acid treatment *in utero* partially rescues the cNCC apoptosis and cleft lip in *Clpex* mutants. Whole mount Nile Blue Sulfate staining of *Clpex* mutants from pregnant dams supplemented with 25ppm folinic acid diet from E0-E9.5 (A,B). Phenotypes observed in *Clpex* mutants from litters treated from E0-E16.5 with control diet (blue), 25ppm folic acid (orange), or 25ppm folinic acid (grey) (C). Summary of the phenotypes of *Clpex* mutants from litters treated with the indicated diets (D). (*p<0.05)

### RNA sequencing reveals changes in patterning genes in *Clpex* mutants

As folinic acid supplementation could not rescue all phenotypes observed in the *Clpex* mutant, we took an unbiased transcriptomic approach to determine the major signaling pathway(s) affected upon reduced *Pgap2* function. RNA sequencing was performed on pooled RNA samples from wild-type and *Clpex* homozygous mutant embryos at E9.5 (5 of each in each RNA pool). Sorting differentially expressed genes in ToppGene showed that the most differentially regulated pathways were sequence-specific DNA binding genes including 39 genes (p=1.79×10^-7^; Table 2; [41]). Among the sequence-specific DNA binding genes, the majority (23/39 genes in the category) were transcription factors which have been implicated in anterior/posterior (A/P) patterning including *Cdx2, Cdx4,Tbxt, Hmx1, Lhx2*, and *Lhx8* [42–49] (Table 3). 13 Anterior patterning genes were statistically significantly downregulated and 10 posterior patterning genes were statistically significantly upregulated (Table 2). We confirmed changes in expression of three of these A/P patterning defects by RNA *in situ* hybridization at E9.5 (Fig. S3). We identified a decrease in *Alx3* in *Clpex* mutants which is both only an anterior patterning gene with a prominent role in frontonasal development and a genetic target of folate signaling [50]. We investigated *Lhx8* because it is expressed in the head at E9.5 and *Lhx8^-/-^* mice develop cleft palate [51]. We found *Lhx8* was decreased in *Clpex* mutant heads. Finally, the master posterior patterning gene *Tbxt (brachyury)* is critical for determining tail length and posterior somite identity [45, 46]. We found *Tbxt* was slightly increased in *Clpex* mutants compared to their overall body size.

**Table 2.**
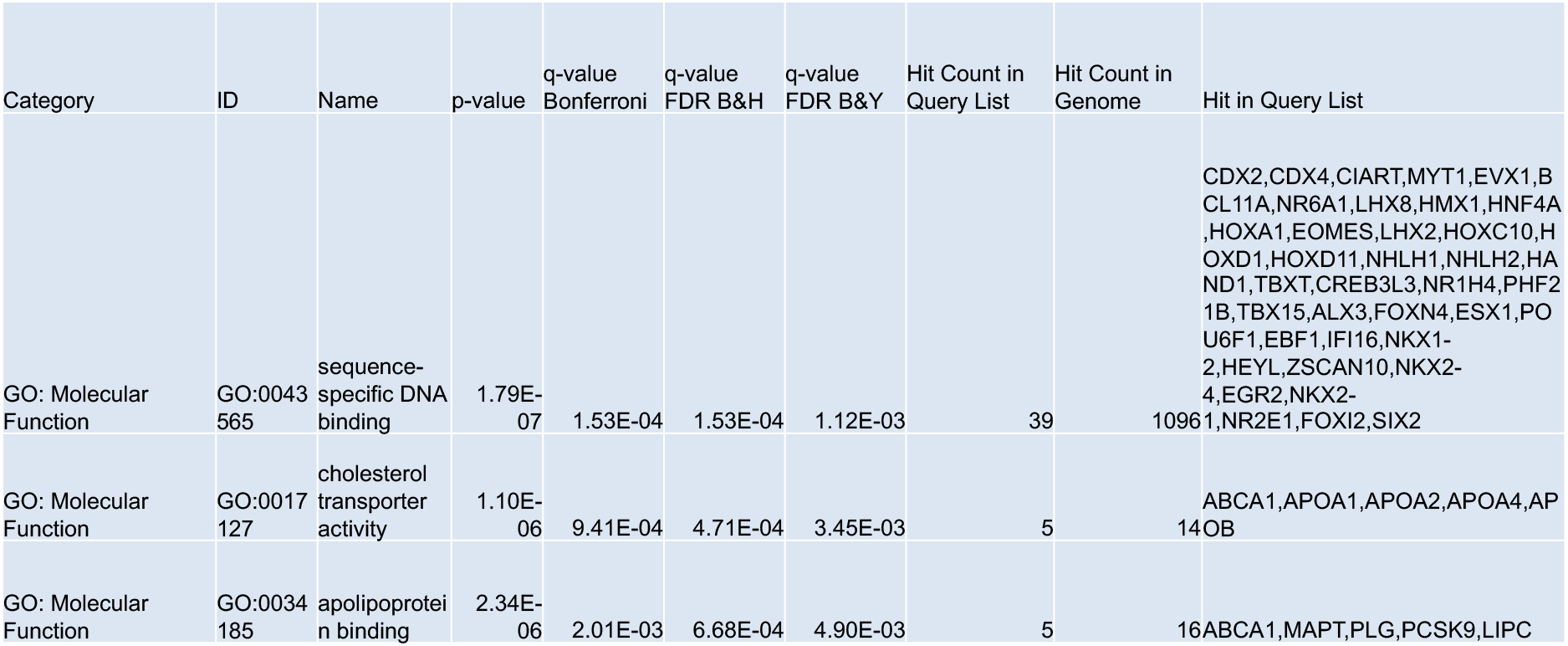
RNA Sequencing ToppGene Pathway Enrichment Analysis.

**Table 3.** Anterior/Posterior Transcription Factors Differentially Expressed in *Clpex* Mutants Compared to Controls.

| A | Anterior TFs |  |  | TPM |  |
| --- | --- | --- | --- | --- | --- |
|  | Gene | Log<sub>2</sub> FC | P-value | WT | *Clpex* |
|  | *Myt1* | -1.09 | 9.63E-09 | 4.84 | 2.05 |
|  | *Lhx8* | -1.56 | 0.004589 | 1.95 | 0.59 |
|  | *Lhx2* | -0.70 | 9.73E-05 | 34.96 | 19.48 |
|  | *Alx3* | -0.71 | 0.001724 | 18.68 | 10.36 |
|  | *Nkx2-4* | -2.37 | 0.009815 | 1.89 | 0.31 |
| B | Posterior TFs |  |  | TPM |  |
|  | Gene | Log<sub>2</sub> FC | P-value | WT | *Clpex* |
|  | *Cdx2* | 1.89 | 0 | 4 | 13.53 |
|  | *Cdx4* | 2.04 | 0 | 3.21 | 12.12 |
|  | *Evx1* | 1.60 | 3.14E-08 | 1.06 | 2.97 |
|  | *Hoxc10* | 1.12 | 8.57E-11 | 10.01 | 19.83 |
|  | *Hoxd11* | 1.07 | 1.69E-06 | 2.43 | 4.59 |
|  | *Nkx1-2* | 1.77 | 2.50E-12 | 1.97 | 6.13 |
|  | *Tbxt* | 1.69 | 0 | 3.35 | 9.89 |

The second and third most altered pathways identified by ToppGene were cholesterol transporter activity (p=1.1×10^-6^), and apolipoprotein binding (p=2.34 x 10^-6^), respectively. Upon closer inspection, the genes in these categories were largely genes expressed in the mesendoderm including *Alpha fetal protein* and the *<u>Apolipoprotein</u>* gene family. We concluded the decreased expression of these genes in the *Clpex* mutant embryos is consistent with a defect in mesendoderm induction, rather than specific cholesterol and apolipoprotein activities.

Collectively, these findings from our transcriptomic analysis suggest other GPI-Aps involved in A/P patterning and mesendoderm development may be affected in *Clpex* homozygous mutant embryos. Multiple other mutations in GPI biosynthesis genes including *Pgap1* and *Pign* lead to defective CRIPTO mediated NODAL/BMP signaling which affects the formation of the A/P axis in the early gastrulating embryo [10, 13, 52, 53]. CRIPTO is a co-receptor for NODAL and necessary for the induction of the anterior visceral endoderm and subsequent forebrain and mesendoderm formation [54, 55]. Mckean et. al. found CRIPTO signaling was impaired in the *Pign^Gonzo^* and *Pgap1^Beaker^* GPI biosynthesis mutants, [10]. Furthermore, stem cells from GPI deficient clones are unable to respond to TGFβ superfamily members due to defects in GPI anchored co-receptor anchoring [10, 52]. Zoltewicz et. al. found mutations in *Pgap1* lead to defective A/P patterning by affecting other major signaling pathways including *Wnt* [53]. Our RNA-Seq results are consistent with the existing literature which has established a critical role for GPI biosynthesis in generating the A/P axis. While this role is well established, few groups have investigated the tissue specific role of GPI biosynthesis after the A-P axis has been established. Interestingly, two studies have found GPI-APs have cell autonomous roles separately in skin and limb development [56, 57]. As we found tissue specific defects in the NCC population in the *Clpex* mutant, we sought to address a larger question and determine the cell autonomous role for GPI-APs in NCC development.

### NCC-specific deletion of *Piga* completely abolishes GPI biosynthesis and leads to median cleft lip, cleft palate, and craniofacial skeletal hypoplasia

We observed cell type specific apoptosis in the cNCCs in the *Clpex* mutant and to further our understanding we sought to determine the cell autonomous role of GPI biosynthesis more generally in these cells. *Phosphatidylinositol glycan anchor biosynthesis, class A (Piga)* is part of the GPI-N-acetylglucosaminyltransferase complex that initiates GPI biosynthesis from phosphatidylinositol and N-acetylglucosamine [1]. *Piga* is totally required for the biosynthesis of all GPI anchors and *Piga* deletion totally abolishes GPI biosynthesis [11, 58, 59]. We first performed RNA whole mount *in situ* hybridization for *Piga* and showed it has a similar regionalized expression as we observed in the *Pgap2* expression experiments. *Piga* expression at E11.5 is enriched in the first branchial arch, heart, limb, and CNS (representative images from n=8 antisense and 2 sense controls over 3 separate experiments) (Fig. 7A-F). However, *Piga* showed a unique enrichment in the medial aspect of both medial nasal processes as opposed to the *Pgap2* expression which appeared to line the nasal pit epithelium (Fig. 7C). Other GPI biosynthesis genes showed a similar regionalization pattern of expression (Fig. S4).

**Figure 7.**
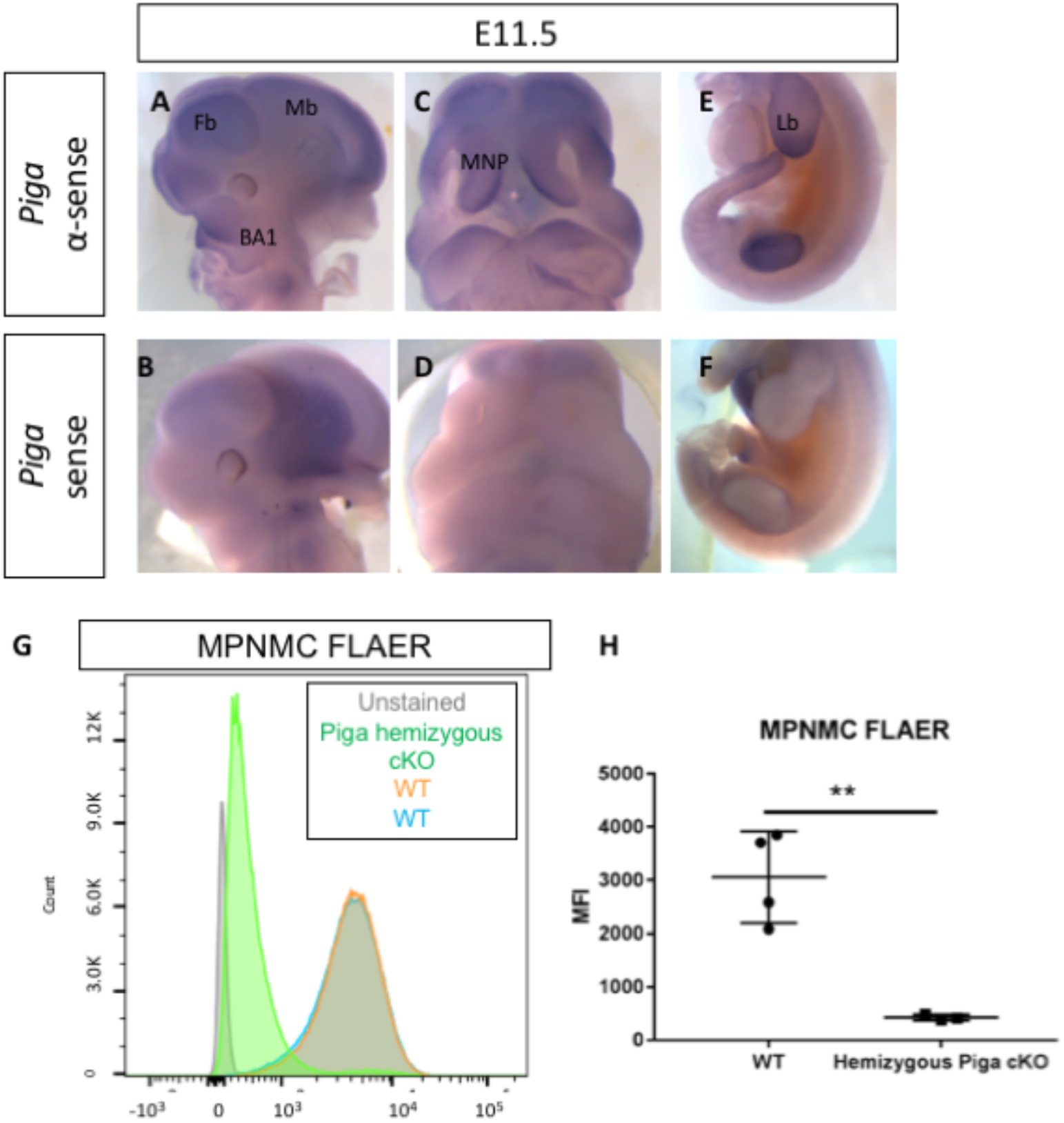
*Piga* expression. *Piga* is expressed in the first branchial arch, medial nasal process, limb bud and deletion of *Piga* in the *Wnt1-Cre* lineage results in NCC cells that lack GPI biosynthesis. WMISH of WT E11.5 embryo stained with αsense *Piga* probe (A, C, E) or sense *Piga* probe (B, D, E). FLAER flow cytometry staining of WT (orange, blue) and *Piga* hemizygous cKO MPNMCs (green). FLAER quantified (H). Fb=Forebrain, Mb=Midbrain, BA1=Branchial Arch 1, MNP=Medial Nasal Process, Lb= Limb bud. **=p<0.001.

To determine the NCC specific role for GPI biosynthesis we generated a novel model of tissue-specific GPI deficiency in the neural crest cell lineage with *Piga^flox/X^; Wnt1-Cre* mosaic conditional KO (cKO) mutants and *Piga^flox/Y^; Wnt1-Cre* hemizygous cKO mutants. To confirm the loss of GPI biosynthesis in these mutants, we cultured Mouse Palatal and Nasal Mesenchymal Cells (MPNMCs) from WT and mutant palates and performed FLAER staining as above. We found mutant MPMNCs lack virtually all GPI anchors on the cell surface (n=4 WT and 3 Mutant cell lines stained in two separate experiments) (Fig. 7G,H).

Analysis of mosaic cKOs at E15.5-16.5 showed mild median cleft lip and cleft palate in all mutants examined (n=6 mutants; Fig. 8A-F). Hemizygous cKOs showed a more severe median cleft lip and cleft palate in all mutants examined (n=7 mutants; Fig. 8G-L). Skeletal preparations to highlight bone and cartilage demonstrates hypoplasia of the craniofacial skeleton, cleft palate (n=5/5 mutants examined; Fig. 8M-R).

**Figure 8.**
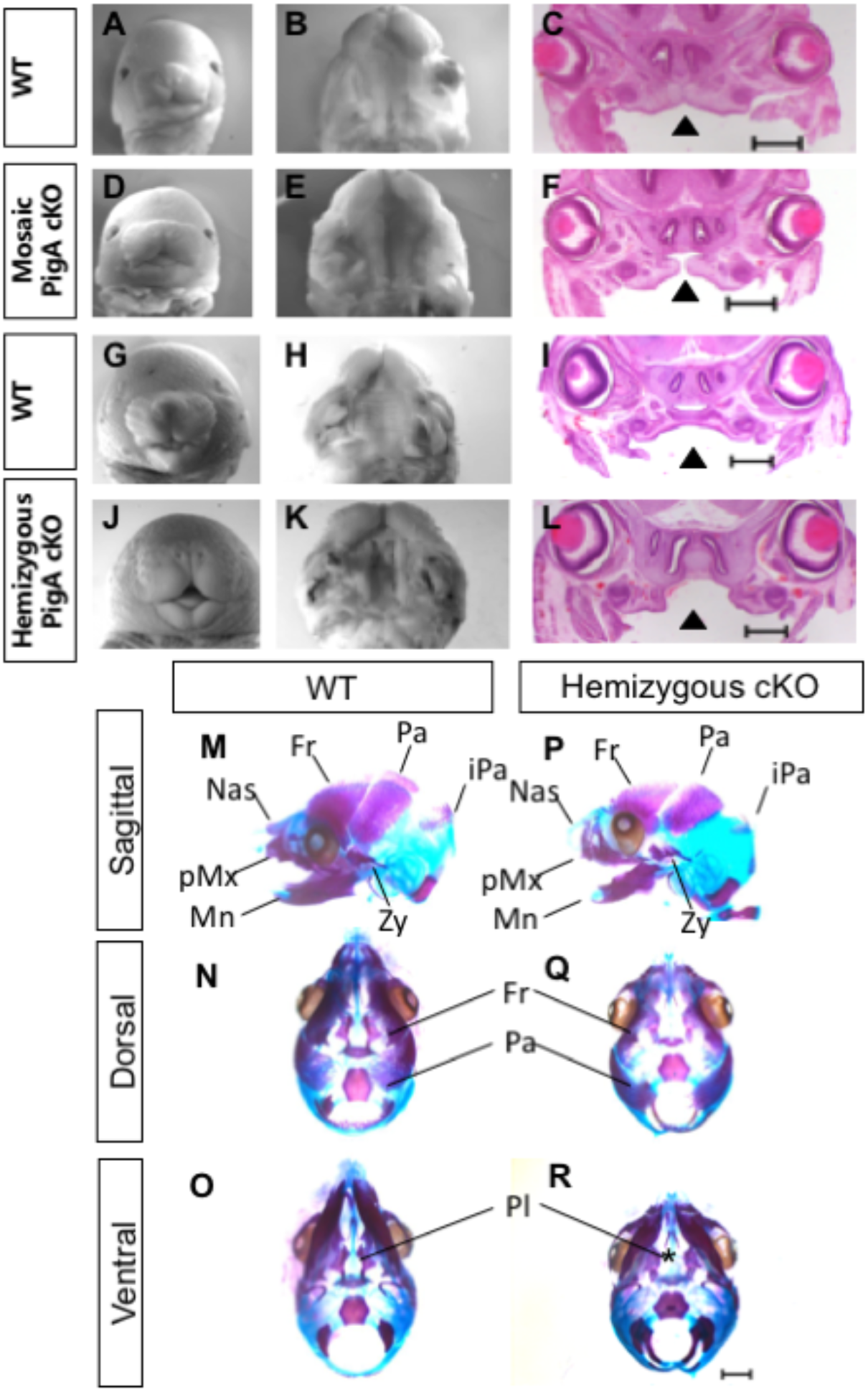
Conditional knockout of *Piga* abolishes GPI biosynthesis in NCCs and leads to craniofacial phenotypes. Whole mount images of E15.5 WT (A), mosaic *Piga* cKO (D), E16.5 WT (G) and hemizygous *Piga* cKO (J). Ventral view of the secondary palate of E15.5 WT (B), Mosaic cKO (E), E16.5 WT (H) and hemizygous cKO (K). H&E staining of E15.5 WT (C), mosaic cKO (F), E16.5 WT (I), and hemizygous cKO (L), arrowhead indicates cleft palate. Alazarin red and alcian blue staining of E16.5 WT skull (M-O) and hemizygous *Piga* cKO skull (P-R). Asterix indicates cleft palate. Fr=Frontal bone, Pa = Parietal bone, iPa= interparietal bone, Zy=Zygomatic bone, Mn=Mandible, pMx= Premaxilla, Nas= Nasal bone. Scale bar indicates 500μM in C, F, I, L and 1mm in M-R.

These data confirm for the first time a cell autonomous role for GPI biosynthesis in cNCCs during development. As we reduced the amount of *Piga* from the mosaic cKO to the hemizgous cKO, we observed a worsening of the cleft lip/palate phenotype including more severe hypoplasia of the palatal shelves and widening of the median cleft lip. These data are consistent with the hypothesis that the dosage of GPI biosynthesis is related to the severity of the phenotype with mutants with less residual GPI anchor expression showing more severe phenotypes. Surprisingly, we found hemizygous cKO mutants are capable of forming all the bones and cartilage of the craniofacial skeleton, though they are all hypoplastic.

These data are consistent with the hypothesis that GPI biosynthesis is involved in the survival of early cNCCs as we observed in the *Clpex* homozygous mutants and not in the later patterning or differentiation of cNCCs. The critical requirement for GPI biosynthesis appears to be at early stages of cNCC survival just after they have migrated from the dorsal neural tube, and before they have committed to differentiation to bone or cartilage.

## Discussion

In this study we aimed to determine the role of GPI biosynthesis in craniofacial development with two novel models of GPI deficiency. First, we characterized the phenotype of the ENU-induced *Clpex* allele which shows partially penetrant cranial neural tube defects, bilateral cleft lip/palate, and edema. We found by mapping and whole exome sequencing the *Clpex* mutation is a homozygous missense allele in the initiating methionine of *Pgap2*, the final enzyme in the GPI biosynthesis pathway. The *Clpex* allele failed to complement a null allele of *Pgap2*, confirming the *Clpex* mutant is caused by a hypomorphic mutation in *Pgap2*. By expression analysis with a *Pgap2^tm1a (lacZ)^* reporter allele, we found *Pgap2* is enriched in the first branchial arch, limb bud, neuroepthelium and around the interior aspect of the nasal pits during lip closure at E9.5-E11.5. In later stages of organogenesis, *Pgap2* is widely expressed and enriched in epithelia. These data argue expression of GPI biosynthesis genes is dynamic during development and not simply uniform and ubiquitous. FLAER flow cytometry and expression of a tagged FOLR1A showed that reduced levels of *Pgap2* affected GPI biosynthesis, though not as severely as a total knockout for the GPI biosynthesis pathway, PIGA^-/-^. Molecular analysis showed the *Clpex* mutants have increased apoptosis in cNCCs and cranial neuroepithelium. Folinic acid diet supplementation *in utero* partially rescued this apoptosis and the cleft lip in *Clpex* mutants. Finally, we generated a NCC tissue specific GPI deficient model to determine the cell autonomous role of GPI biosynthesis. *Piga^flox/X^; Wnt1-Cre* mosaic cKO mutants and *Piga^flox/Y^; Wnt1-Cre* hemizygous cKO mutants displayed fully penetrant median cleft lip/cleft palate and craniofacial hypoplasia similar to our germline *Clpex* mutant, confirming a cell autonomous role for GPI biosynthesis in craniofacial development.

Contrary to previous studies of other GPI biosynthesis pathway genes, we found *Pgap2* clearly shows enriched expression in certain tissues during certain stages of development. We observed a similar pattern in *Piga* RNA expression suggesting GPI biosynthesis genes share similar gene enrichment domains. These tissues are the most affected in GPI biosynthesis mouse mutants and include the craniofacial complex, CNS, limb, and heart. This may mean *Pgap2* and other GPI biosynthesis genes are required in certain tissues for anchoring GPI-APs critical to that tissue. Alternatively, these areas may be particularly “GPI-rich.”

A variety of mutants have been described in the GPI biosynthesis pathway with a wide array of phenotypes [1, 4]. While germline mutants in this pathway remain poorly understood, recent research in Paroxysmal Nocturnal Hemoglobinuria (PNH) caused by somatic mutations in *PIGA* has revolutionized our understanding of GPI deficiency related pathology. In PNH, clones of GPI deficient hematopoietic stem cells proliferate in the bone marrow and give rise to blood cells that lack GPI-anchored CD55/59 which are required to prevent complement-mediated lysis of red blood cells. PNH patients suffer from episodes of hemolysis and thrombosis which can be deadly [60]. Blockade of complement in these patients via eculizumab, a monoclonal antibody that inhibits the conversion of C5 to C5a, has been shown to greatly improve survival [61–64]. Thus, a single GPI-AP seems to be largely responsible for the disease observed in these patients.

In this study we aimed to identify a single GPI-AP that could be responsible for all the phenotypes observed in our germline GPI biosynthesis *Clpex* mutant. Of the known GPI-AP knockout models, *Clpex* shares the most phenotypic overlap with the *Folr1^-/-^* mouse. We directly tested the hypothesis that FOLR1 deficiency is solely responsible for the *Clpex* phenotype by dietary supplementation of folinic acid during embryonic development. To our surprise, folinic acid supplementation could partially rescue the cleft lip phenotype but not the NTD or cleft palate (Fig. 6C,D). These data also argue high dose folinic acid *in utero* may be a possible therapeutic for some phenotypes in patients with GPI biosynthesis variants. Further research is required to test whether the positive effect of folinic acid on the *Clpex* mutants could be observed in other GPI biosynthesis mutants. These data also argue the phenotypes observed in germline *Clpex* mutants do not share a single mechanism and are not due to the loss of a single GPI-AP. Given the varied response in different tissues to the rescue regimens utilized here, it is likely loss of different GPI-

Many GPI-APs could be responsible for the phenotypes we observe in the *Clpex* mutant but were not tested explicitly in this work. Notably, the two receptors for Glial Derived Neurotrophic Factor (GDNF) are GPI-anchored (GFRA1, GFRA2). GFRA1 and GFRA2 are known to be critical for the survival and development of NCCs in the gut during enteric nervous system development [65–67] Interestingly, *Gfra1and 2* are expressed in the craniofacial complex during development [68, 69]. Whether GDNF plays a crucial in cNCC survival remains to be explored. Other candidate GPI-APs that may be affected by loss of *Pgap2* include one form of Neural Cell Adhesion Molecule (NCAM), a critical neural cell adhesion. NCAM^-/-^ mice display defects in neural tube development including kinking and delayed closure [70]. A third candidate includes the glypican family members which are GPI anchored heparin sulfate proteoglycans that play critical roles in cell-cell signaling and have been shown to modulate critical patterning gradients in the neural tube and face including Sonic Hedgehog and Wnt [71–74]. Other GPI-AP knockout models display NTDs including *Repulsive guidance molecule A/B (Rgma)* and *Ephrin A5 (Efna5). However, Rgma^-/-^* mice do not develop increased apoptosis in the neuroepithelium as observed in *Clpex* mutants [75]. *Efna5^-/-^* mice appear to form DLHPs though the neural folds do not fuse in the midline which is less severe than the defect we observe in *Clpex* mutants [22]. Therefore, we find it unlikely the loss of these GPI-APs are primarily responsible for the defects observed in the *Clpex* mutant although contributions to the phenotype may come from abnormal presentation of one or several of these GPI-Aps on the cellular membranes.

It has been known for decades that treatment of embryos with phospholipase C to release GPI-APs from the cell surface causes NTD *in utero* [76]. To investigate the cause of the NTD in *Clpex* mutants, we performed histological and immunohistochemical analysis of the mutant at neurulation stages. We found the *Clpex* mutant fails to form dorsolateral hinge points and the cranial neuroepithelium is apoptotic in the region of the developing DLHP. Neuroepithelial apoptosis was restricted to the midbrain/hindbrain boundary and likely explains why *Clpex* mutants develop cranial NTDs as opposed to caudal NTDs such as spina bifida. These cellular defects likely underlie the NTD but the cause of the neuroepithelial apoptosis remains unclear as the NTD did not respond to folinic acid supplementation. It remains controversial, but the NTD in *Folr1^-/-^* mice may be related to an expansion of the *Shh* signaling domain that patterns the neural tube [36, 77]. Indeed, many *Shh* gain-of-function mutants develop NTD as *Shh* expansion impairs the formation of DLHPs and closure of the neural tube [77]. Our RNA sequencing analysis did not identify a dysregulation in the *Shh* signaling pathway so there are likely differences in the mechanism responsible for the NTD in *Folr1^-/-^* mice and *Clpex* mutants.

To determine alternative mechanisms responsible for the *Clpex* phenotype, we performed RNA sequencing from E9.5 WT and *Clpex* mutants. We found the largest differences in gene expression were in A/P patterning genes and mesendoderm induction genes. The A/P axis and induction of mesendoderm has been shown to require GPI-anchored CRIPTO, a Tgfβ superfamily member co-receptor of NODAL. A variety of studies have shown CRIPTO/Tgfβsuper family members pathway function is impaired in GPI biosynthesis mutants because CRIPTO is GPI-anchored and cleavage of the anchor affects CRIPTO function [10, 52].

While GPI deficiency has been studied in the context of A/P patterning, this is the first study to implicate GPI biosynthesis in the survival of neural crest cells in a cell autonomous fashion. Indeed, the enrichment of *Piga* in the developing medial nasal process and the median cleft lip/cleft palate and craniofacial hypoplasia in our *Pig*a cKOs confirms a unique cell autonomous role for GPI biosynthesis in these structures. Interestingly, these mutants do not show a complete loss of the craniofacial skeleton, rather a general, mild hypoplasia consistent with a role for GPI biosynthesis in early NCC survival, but not later patterning or differentiation.

Our study provides potential mechanistic explanations for the developmental defects observed in a GPI biosynthesis mutant model. We propose GPI biosynthesis is involved in anchoring critical survival factors for NCCs and the neuroepithelium. In GPI deficient states, NCCs undergo apoptosis leading to hypoplastic nasal processes and palatal shelves. As we reduced the degree of GPI biosynthesis from the germline *Clpex* mutant hypomorph to our totally GPI deficient NCC cKO model, we observed a worsening of the craniofacial phenotype as witnessed by the fully penetrant cleft lip/cleft palate and craniofacial hypoplasia. These data argue the degree of GPI deficiency correlates with the severity of the phenotype. In the neuroepithelium, loss of neuroepithelial cells at the DLHPs result in failure to bend and close the neural tube. Conditional ablations of critical

GPI biosynthesis genes in other affected tissues including the CNS and heart will likely lead to new understandings of the diverse pathology of inherited glycophosphatidylinositol deficiency.

## Materials and Methods

### Animal husbandry

All animals were maintained through a protocol approved by the Cincinnati Children’s Hospital Medical Center IACUC committee (IACUC2016-0098). Mice were housed in a vivarium with a 12-h light cycle with food and water *ad libitum*. The Clpex line was previously published by Stottmann et. al. [9]. *Piga^flox^* (*B6.129-Piga^tm1^*) mice were obtained from RIKEN and previous were previously generated by Taroh Kinoshita and Junji Takeda [11]. *Wnt1-Cre* (B6.Cg-*H2afv*^*Tg(Wnt1-cre)11RthTg*(Wnt1-GAL4)11Rth^/J) mice and R26R LacZ reporter (B6.129S4 *Gt(ROSA)26Sor^tm1Sor^/J;* R26R^Tg^) mice were purchased from Jackson Laboratories and previously published. *Pgap2^null^ (Pgap2^tm1a(EUCOMM)Wtsi^)* mice were obtained from EUCOMM and genotyped using their suggested primers. Primers used to genotype all animals are listed in Table S1. Sample2Snp custom Taqman probes were designed by Thermo-Fisher and used to genotype the point mutation in the *Clpex* line.

### Mapping and Sequencing

Mapping of the *Clpex* mutation was previously described [9]. Whole exome sequencing was done at the CCHMC DNA Sequencing and Genotyping Core. The *Pgap2* exon 3 variant was Sanger sequenced using the Zymo DNA clean & Concentrator kit (Zymo Research Corporation, Irvine, CA).

### Whole Mount *In Situ* Hybridization

RNA *in situ* hybridization was performed as previously described [78]. Briefly, whole E8-E11.5 embryos were fixed overnight in 4% PFA at 4°C and dehydrated through a methanol series. Samples were treated with 4.5μg/mL Proteinase K for 7-13 minutes at room temperature, post-fixed in 4% PFA/0.2% glutaraldehyde and blocked with hybridization buffer prior to hybridization overnight at 65°C with constant agitation. The samples were washed and incubated with an anti-Dig antibody (Roche #11093274910) o/n at 4°C. Embryos were washed and incubated with NBT/BCIP (SIGMA) or BM Purple (Roche #11442074001) from 4 hours at room temperature to o/n at 4°C.

*Piga* (#MR222212), *Pgap2* (#MR2031890) *Pigp* (#MR216742), *Pigu* (#MR223670), *Pigx* (#MR201059), *Lhx8* (#MR226908), and *Tbxt* (#MR223752) plasmids were obtained from Origene (Rockville, MD). Antisense probes were generated from PCR products containing T3 overhangs. *Piga, Pgap2, Pigp, Pigu, Pigx, Lhx8*, and *Tbxt* antisense probes were generated from 910, 750, 556, 952, and 416, 519, and 665 base pair products, respectively. The PCR products were purified, *in vitro* transcription was performed with digoxigenin-labeled dUTP (Roche #11277073910), and the probe was purified with the MEGAclear Transcription Clean-up kit (Thermo #AM1908) per the manufacturer’s instructions. For sense probes, the plasmids were cut with XhoI restriction enzyme after the coding sequence and T7 RNA polymerase was used for *in vitro* transcription. The *Alx3* probe was generated by *in vitro* transcription of a 790bp PCR product from Alx3 plasmid (DNASU #MmCD00081160) containing T3 overhangs.

### MEF/MPNMC production and FLAER staining

MEFs were generated from E13.5 embryos. Embryos were dissected in PBS, decapitated, and eviscerated. The remaining tissue was incubated in trypsin o/n at 4°C to allow for enzymatic action on the tissue and remaining fibroblasts were passaged in complete DMEM containing 10%FBS and penicillin/streptomycin. MEFs were stained within three passages of their isolation. MEFs and 293T cells were stained with 5μL of Alexafluor-488 proaerolysin (FLAER)/1×10^6^ cells (CedarLane Labs, Burlington, Ontario, Canada) and flow cytometry was performed on Becton-Dickinson FACSCanto II flow cytometer in the CCHMC Research flow cytometry core. Mouse Palatal Nasal Mesenchymal Cells (MPNMCs) were generated from E13.5-E14.5 microdissected embryo heads in a protocol similar to that used for MEPMS [79]. The lower jaw, eyes and brain were removed and the remaining upper jaw and nasal mesenchyme were lysed in 0.25% trypsin for 10 min at 37°C, passaged through a P1000 pipette several times to create a single cell suspension, and cultured in 12 well plates. These cells displayed a stellate mesenchymal cell appearance after culture overnight. They were then stained after 72 hours from isolation with FLAER.

### CRISPR Knockout/Knock-in gene editing

We utilized a double guide approach to generate knockout clones with deletions in *PGAP2* and *PIGA* in HEK293T cells. Two small guide RNAs targeting exon 3 of either *PGAP2* or *PIGA* were designed using Benchling software (Benchling, San Francisco, CA) and 5’ overhangs were added for cloning into CRISPR/Cas9 PX459M2 puromycin-resistance vector [80]. We also generated a single gRNA and donor oligonucleotide for homologous recombination to recapitulate the *Clpex* mutation in 293T cells (Integrated DNA Technologies ultramer). We cloned these guides into the PX459M2 plasmid using the one-step digestion-ligation with BbsI enzyme as described by Ran et. al. [81]. Two guides per gene were transfected in WT 293T cells using Lipofectamine 3000 and cells were selected for transfection by 3 days of culture in 10μg/mL puromycin. Transfected cells were plated at clonal density into a 96 well plate and single clones were scored approximately one week post seeding. Single clones were Sanger sequenced to confirm deletion of the target exon 3 sequence of either *PIGA* or *PGAP2. PIGA* clones carry a 50bp out-of-frame deletion in *PIGA* and lack virtually all GPI expression on the cell surface by FLAER flow cytometry staining. *PGAP2* clones carry a 121bp out-of-frame deletion in *PGAP2*. Clpex Knock-in clones were Sanger sequenced to identify clones carrying the desired knock-in mutation and clones with indels were discarded. Primers for sgRNA cloning and PCR amplification of targeted regions can be found in Table S1. Sequencing of clones is presented in Fig. S1.

### Immunofluorescence

293T cells were transfected with FOLR1-myc constructs (Origene #RC212291using Lipofectamine 3000, incubated for 48 hours, then fixed for 15 min in 4% PFA, and blocked in 4% normal goat serum. They were stained o/n at 4°C with 1:500 rabbit anti-myc (Abcam ab9106), washed the next day and stained with 1:500 488-congugated goat anti-rabbit (Thermo #A11008). They were then stained for 5 min with 5μg/mL wheat germ agglutinin (Thermo Fischer #W21405) and counter-stained with DAPI. They were visualized on a Zeiss widefield microscope

E9.5 embryos were dissected, fixed in 4% PFA o/n, equilibrated in 30% sucrose o/n, cryo-embedded in OCT, and sectioned from 10-20μM by cryostat. Sections were subjected to antigen retrieval by citrate retrieval buffer, blocked in 4% normal goat serum, incubated in primary antibody 1:20 mouse anti-AP2 (Developmental Studies Hybridoma Bank, University of Iowa, 3B5 supernatant) and 1:300 rabbit anti-Cleaved Caspase 3 (Cell Signaling Technology, Danvers, MA #9661) o/n in humid chamber. Sections were incubated with secondary antibody 1:1000 Alexafluor 488-congugated goat anti-rabbit (Thermo #A11008) and 1:1000 Alexafluor 594 conjugated goat anti-mouse (Thermo A11008) and counterstained with DAPI. Sections were imaged on Nikon C2 confocal microscope and CC3+ cells and AP2+ cells were quantified with Nikon Elements software brightspot analysis.

### Western blotting

293T cells were transfected with FOLR1-myc constructs using Lipofectamine 3000, incubated for 48 hours and lysed in RIPA buffer containing Protease Inhibitor cocktail (Roche #11697498001). Lysate protein concentration was determined by BCA assay and electrophoresis was performed on a 10% Tris-glycine gel. Protein was transferred to a PVDF membrane, blocked in Odyssey blocking buffer and incubated o/n at 4°C with 1:1000 Rabbit anti-myc (abcam ab9106) and 1:1000 Mouse anti-Tubulin (Sigma #T6199) antibodies). Membranes were washed and incubated for 1 hour in 1:15000 goat antirabbit IRDye 800CW (LICOR # 926-32211)and 1:15000 goat anti-mouse IRDye 680Rd (LICOR, #926-68070) and visualized on LICOR Odyssey imaging system.

### Nile Blue Sulfate Viability Stain

E9.5 embryos were dissected in PBS and placed immediately into 10ml of complete DMEM containing 50ul of 1.5% Nile blue stock solution (Sigma #N5632-25G) for 30 minutes at 37°C to allow for uptake of the dye. They were then washed for 1 hour in PBS at 4°C and immediately visualized.

### Histology

Whole embryos E8-E16.5 were fixed in formalin and embedded in paraffin for coronal sectioning and stained with hematoxylin and eosin using standard methods.

### NCC lineage trace and Xgal staining

*Clpex* heterozygous females were crossed to *Wnt1-Cre* R26R transgenic mice as described in results. Whole embryos were fixed in 4% PFA for 15 minutes at RT, washed in lacZ buffer, and stained in a solution containing 1mg/mL X-gal (Sigma #B4252) [82]. They were washed 3 times in PBS-T and imaged after several hours in X-gal stain at room temperature.

### Diet

*Clpex* pregnant dams were treated with either control chow, chow + 25ppm folic acid, or chow + 25ppm folinic acid generated by Envigo (Indianapolis, Indiana) from E0-E16.5 *ad libitum*. They were euthanized at either E9.5 or E16.5 to assess phenotype

### RNA Sequencing

5 WT and 5 *Clpex* mutant E9.5 embryos were snap frozen on dry ice. RNA was isolated and pooled samples of each genotype were used for paired-end bulk-RNA sequencing (BGI-Americas, Cambridge, MA). RNA-Seq analysis pipeline steps were performed using CSBB [Computational Suite for Bioinformaticians and Biologists: https://github.com/csbbcompbio/CSBB-v3.0]. CSBB has multiple modules, RNA-Seq module is focused on carrying out analysis steps on sequencing data, which comprises of quality check, alignment, quantification and generating mapped read visualization files. Quality check of the sequencing reads was performed using FASTQC (http://www.bioinformatics.bbsrc.ac.uk/projects/fastqc). RNA-Seq reads for the mutant and wildtype were paired-end and had ~43 and ~31 million reads respectively. Reads were mapped (to mm10 version of Mouse genome) and quantified using RSEM-v1.3.0 [83]. Differential expression analysis was carried out by EBSeq [https://www.biostat.wisc.edu/~kendzior/EBSEQ/] [84]. Differential transcripts are filtered based on LogFC and p-value. Filtered DE transcripts are used for functional and pathway enrichment using toppgene [https://toppgene.cchmc.org/] [41].

### Skeletal Preparation

For skeletal preparation, E16.5-E18.5 embryos were eviscerated and fixed for 2 days in 95% ethanol. They were stained overnight at room temperature in Alcian blue solution (Sigma #A3157) containing 20% glacial acetic acid. They were destained for 24 hours in 95% ethanol and slightly cleared in a 1% KOH solution o/n at room temperature. They were then stained o/n in Alazarin red solution (Sigma #A5533) containing 1% KOH. They were then cleared for 24 hours in 20% glycerol/1%KOH solution. Finally, they were transferred to 50% glycerol/50% ethanol for photographing.

### Barrier Function assay

E18.5 embryos were dehydrated through a methanol series and then rehydrated. Next, they were placed in 0.1% Toludine Blue (Sigma #89640) in water for 2 minutes on ice. They were destained in PBS on ice and imaged.

### Statistical Analysis

Statistical analysis was performed using Graphpad Prism (GraphPad Software, San Diego, CA). All tests but diet results were unpaired, two-tailed t-tests and significance was labelled with one asterix=p<0.05 and two asterix= p<0.001. For statistical analysis of phenotypes observed for embryos under varying diet conditions, z-test of proportions was used.

## ACKNOWLEDGEMENTS

This works was supported by the Cincinnati Children’s Research Foundation, NIH (R.W.S. R01NS085023) and the American Cleft Palate - Craniofacial Association (Paul W. Black, MD Grant for Emerging Researchers to M.J.L.).

## AUTHOR CONTRIBUTIONS

M.L., T.R., P.C., R.W.S. generated and analyzed the data. M.L. and R.W.S. conceived, designed, and wrote the study.

