## Supplemental Figure 1 for "Glycosylphosphatidylinositol Biosynthesis and Remodeling are Required for Neural Crest Cell, Cardiac and Neural Development"

Lukacs_Fig S1

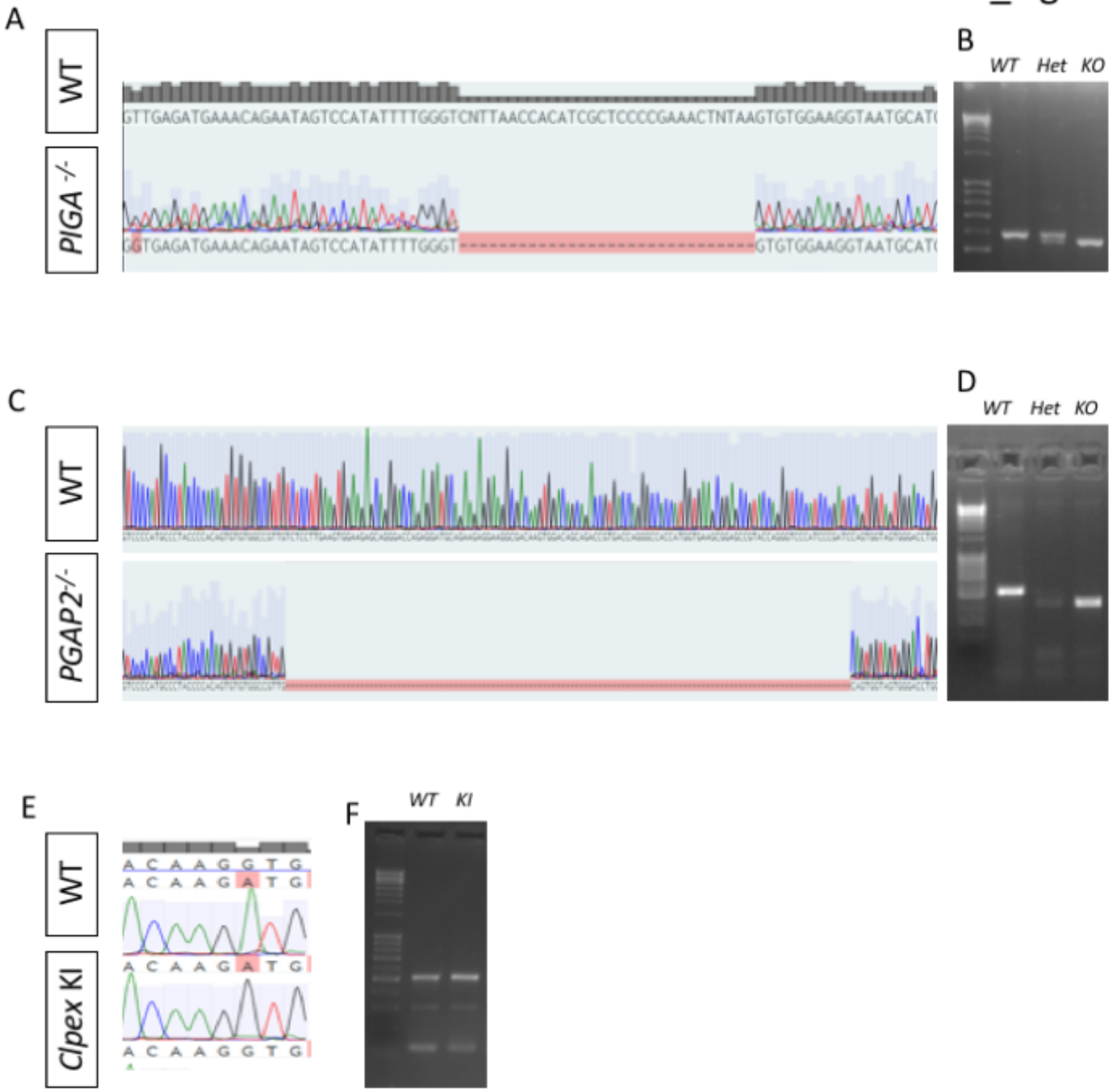


**Supplementary Figure 1. Sequencing of CRISPR/Cas9 generated *PIGA*$^{-/-}$, *PGAP2*$^{-/-}$, and *Clpex* KI 293T clones.** WT human sequence and Sanger Sequencing of *PIGA*$^{-/-}$ clone showing a 29bp deletion in PIGA exon 3 (A). PCR of exon 3 of PIGA showing a heterozygous clone with a small deletion and the KO with the 29bp deletion (B). Sanger Sequencing of WT 293T *PGAP2* exon 3 and *PGAP2*$^{-/-}$ clone showing a 121bp deletion (C). PCR of *PGAP2* exon 3 showing WT 293T, heterozygous clone with two deletions and the *PGAP2*$^{-/-}$ clone with a single large deletion (D). Sanger Sequencing of *PGAP2* exon 3 in WT 293T and *Clpex* Knock-in (KI) clone with the highlighted A>G mutation (E). PCR of PGAP2 exon 3 with WT 293T and Clpex KI clone DNA (F).
