## Supplemental Figure 2 for "Glycosylphosphatidylinositol Biosynthesis and Remodeling are Required for Neural Crest Cell, Cardiac and Neural Development"

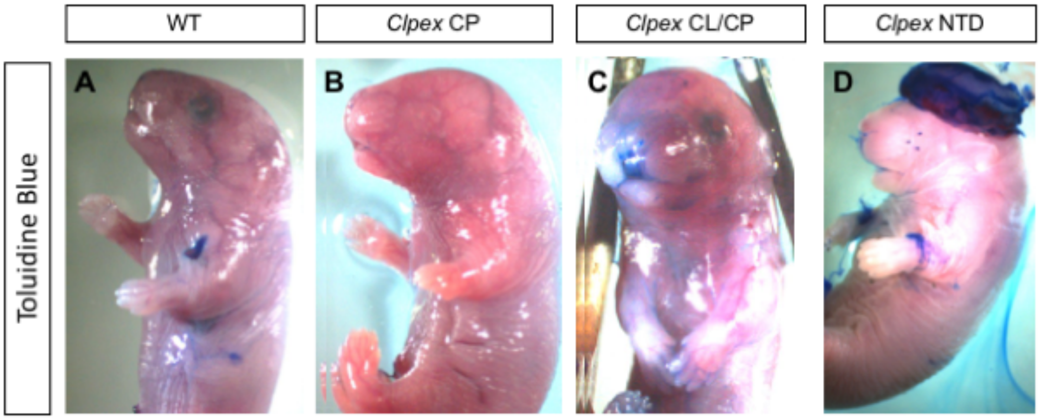


**Supplementary Figure 2. *Clpex* mutants do not display a defect in barrier formation**. E18.5 WT (A), *Clpex* cleft palate mutant (B), *Clpex* cleft lip/cleft palate mutant (C), and *Clpex* neural tube defect mutant (D) stained with Toludine Blue.
