## Supplemental Figure31 for "Glycosylphosphatidylinositol Biosynthesis and Remodeling are Required for Neural Crest Cell, Cardiac and Neural Development"

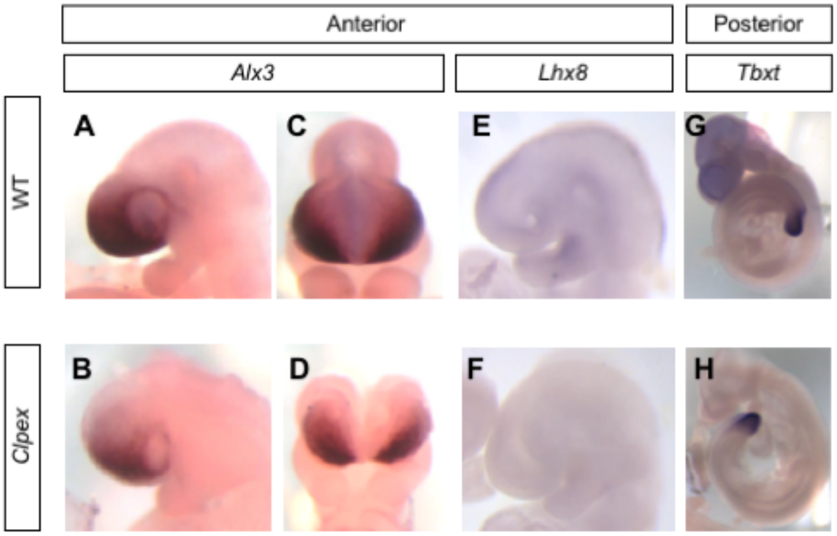


**Supplementary Figure 3. *Clpex* mutants display defects in expression of anterior/posterior patterning genes.** E9.5 WT (A,C) and *Clpex* mutant (B, D) WMISH with $\alpha$*Alx3* probe, an anterior pattering gene. E9.5 WT (E) and *Clpex* mutant (F) WMISH with $\alpha$*Lhx8* probe, an anterior patterning gene. E9.5 WT (G) and Clpex mutant (H) WMISH with $\alpha$*Tbxt* (*Brachyury*) probe, a posterior patterning gene.
