## Supplemental Figure 4 for "Glycosylphosphatidylinositol Biosynthesis and Remodeling are Required for Neural Crest Cell, Cardiac and Neural Development"

Lukacs_Fig S4

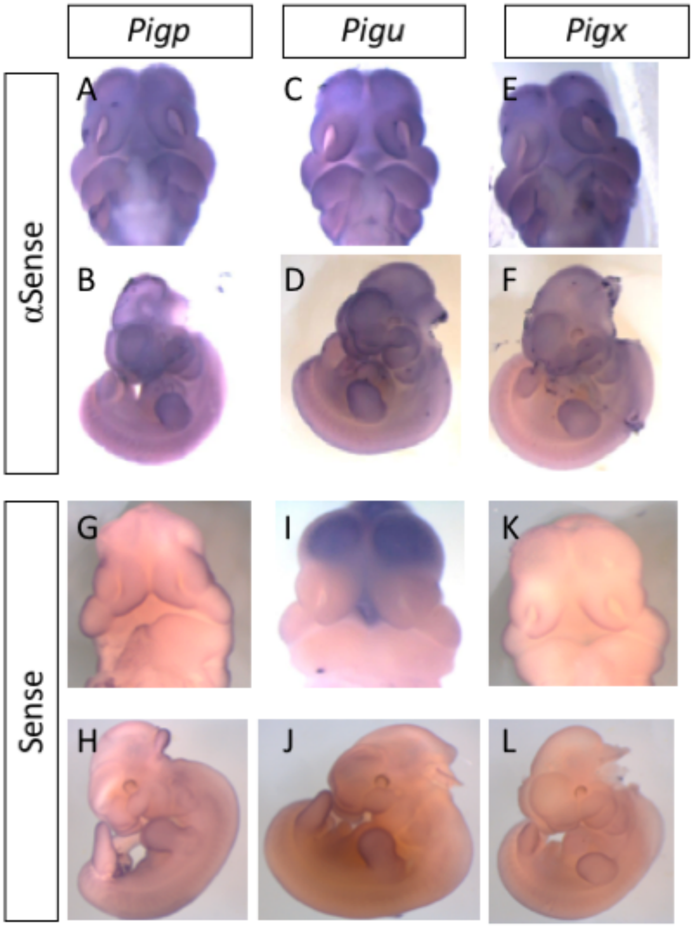


**Supplementary Figure 4. GPI biosynthesis genes show increased expression in the first branchial arch, limb bud, and forebrain.** WT E11.5 RNA *in situ* hybridization with *Pigp* antisense (A,B) and sense (G,H); *Pigu* antisense (C,D) and sense (I,J) and *Pigx* antisense (E,F) and sense (K,L) probes.
