## Supplemental Table 1 for "Glycosylphosphatidylinositol Biosynthesis and Remodeling are Required for Neural Crest Cell, Cardiac and Neural Development"

**Table S1. Primers. Primers are listed by name, sequence, and purpose according to t**

| Primer | Sequence |
|---|---|
| mPiga G4 | ACCTCCAAAGACTGAGCTGTTG |
| mPiga G3 | CCTGCCTTAGTCTTCCCAGTAC |
| mPigalox | TGTGGGTTTCAGTTCATTTCAGA |
| Clpex S2S F | ACAACACACCTTCCATCCTTAGG |
| Clpex S2S R | CATCCCGGTCCAATGTCAGT |
| Clpex S2S Reporter sequence 1 | ACCTGGTACATCTTGTC |
| Clpex S2S Reporter sequence 2 | CCTGGTACACCTTGTC |
| hPIGA exon 3 F | CGCATGCAGTTAAAACCAAA |
| hPIGA exon 3 R | GGGAAAAGACCCAGATCTCC |
| hPGAP2 exon3L F | CAGGGCAAGAGCTATCCAAG |
| hPGAP2 exon3L R | ACCCTCTAGCCCAATCCTGT |
| mPiga WMISH F | TGTCACCCATGCTTATGGAA |
| mPiga WMISH R | ATTAACCCTCACTAAAGGCAATGTCCCCGACTTCACTT |
| WelTrust CAS_R1_Term | TCGTGGTATCGTTATGCGCC |
| WelTrust Pgap2_F9 | GGGCTCAGGAGTACAAGCTG |
| IMR1084 | GCG GTC TGG CAG TAA AAA CTA TC |
| IMR1085 | GTG AAA CAG CAT TGC TGT CAC TT |
| hPIGA exon 3 Guide 1 F | CACCGGATATTTCTGACAGAGTTC |
| hPIGA exon 3 Guide 1 R | AAACGAACTCTGTCAGAAATATCC |
| hPIGA exon 3 Guide 2 F | CACCGAATTTCATAATTGGAGGAGA |
| hPIGA exon 3 Guide 2 R | AAACTCTCCTCCAATTATGAAATT |
| Pgap2 Guide 4 F | caccgCTTCCACTTCAAGGAGACAA |
| Pgap2 Guide 4 R | aaacTTGTCTCCTTGAAGTGGAAGc |
| Pgap2 Guide 6 F | caccgAGGGTCCCATCCCGATCCAG |
| Pgap2 Guide 6 R | aaacCTGGATCGGGATGGGACCCTc |
| Clpex KI sgRNA F | caccg CAGGTCTGACAAGATGTACC |
| Clpex KI sgRNA R | aaac GGTACATCTTGTCAGACCTGc |
| Clpex KI clone donor oligo | G*G*CTGCCTGGGGCCCTGACAGCATGCCACTCCATCCCCAGGT<br>CTGACAAGgTGTACCAAGTCCCACTACCACTGGATCGGGATGGG<br>ACCCTGGTACGGCTC*C*G |
| Kdm5 intron F | TGAAGCTTTTGGCTTTGAG |
| Kdm5 intron R | CCGCTGCCAAATTCTTTGG |
| mAlx3 in situ F | TGGAACCCTACCTCCCAGAG |
| mAlx3 in situ R | ATTAACCCTCACTAAAGGTAGTCACCATCCGGAGAAGG |
| Brachyury WMISH F | CCGGTGCTGAAGGTAAATGT |
| Brachyury WMISH R | ATTAACCCTCACTAAAGGTGACCGGTGGTTCCTTAGAG |
| mLHX8 WMISH F | TGTAAGCTGGAGGGAAAGGA |
| mLHX8 WMISH R | ATTAACCCTCACTAAAGGTTGGATGATTGACGTCTTGC |
| PigP WMISH F | GCAGGGTCTGCACTCACC |
| PigP WMISH R | ATTAACCCTCACTAAAGGGGTATTGAGTTCTTTGGCTCCT |
| PigU WMISH F | GCACTGTTGGATCTGGGAGT |
| PigU WMISH R | ATTAACCCTCACTAAAGGAGCAGACGATGATGATGCAG |
| PigX WMISH F | AAGCCCCCAACTACTTGTCC |
| PigX WMISH R | ATTAACCCTCACTAAAGGTGGCCATATTTGAAAACAGC |
| Pgap2 in situ probe primer F | TTATAAAAGCTTGCGGCCGCAGAATATTGTACCAGGTCCCACTGA |

| | |
|---|---|
| Pgap2 in situ probe primer R | GCTCTAGAAATTAACCCTCACTAAAGGCAGCTCTTTGTTCCCGAAG[illegible] |
| Pgap2 Sanger F | TGGCAGAGAGTTGCAGAGAA |
| Pgap2 Sanger R | AGCCCATGAGAACTGGCTAA |

heir role in this work.

| Purpose |
| --- |
| Genotyping of Piga$^{flox}$ mice |
| Genotyping of Piga$^{flox}$ mice |
| Genotyping of Piga$^{flox}$ mice |
| Genotyping of Clpex colony |
| Genotyping of Clpex colony |
| Genotyping of Clpex colony |
| Genotyping of Clpex colony |
| Genotying PIGA$^{-/-}$ 293T clones |
| Genotying PIGA$^{-/-}$ 293T clones |
| Genotying PGAP2$^{-/-}$ and Clpex KI 293T clones |
| Genotying PGAP2$^{-/-}$ and Clpex KI 293T clones |
| Generation of Piga antisense RNA probe from plasmid |
| Generation of Piga antisense RNA probe from plasmid |
| Genotyping of Pgap2$^{null/LacZ}$ mice |
| Genotyping of Pgap2$^{null/LacZ}$ mice |
| Genotyping of Wnt1-Cre mice |
| Genotyping of Wnt1-Cre mice |
| Cloning of PIGA sgRNA into PX459 Cas9 plasmid |
| Cloning of PIGA sgRNA into PX459 Cas9 plasmid |
| Cloning of PIGA sgRNA into PX459 Cas9 plasmid |
| Cloning of PIGA sgRNA into PX459 Cas9 plasmid |
| Cloning of PGAP2 sgRNA into PX459 Cas9 plasmid |
| Cloning of PGAP2 sgRNA into PX459 Cas9 plasmid |
| Cloning of PGAP2 sgRNA into PX459 Cas9 plasmid |
| Cloning of PGAP2 sgRNA into PX459 Cas9 plasmid |
| Cloning of PGAP2 sgRNA into PX459 Cas9 plasmid to generate Clpex KI clones |
| Cloning of PGAP2 sgRNA into PX459 Cas9 plasmid to generate Clpex KI clones |
| Donor for homologous repair when used in combination with Clpex KI sgRNA to recapitulate the Clpex mutation in 293T ce |
| Genotyping of male/female mice using sexually dimprohic KDM5 intronic locus |
| Genotyping of male/female mice using sexually dimprohic KDM5 intronic locus |
| Generation of Alx3 antisense RNA probe from plasmid |
| Generation of Alx3 antisense RNA probe from plasmid |
| Generation of Tbxt antisense RNA probe from plasmid |
| Generation of Tbxt antisense RNA probe from plasmid |
| Generation of Lhx8 antisense RNA probe from plasmid |
| Generation of Lhx8 antisense RNA probe from plasmid |
| Generation of Pigp antisense RNA probe from plasmid |
| Generation of Pigp antisense RNA probe from plasmid |
| Generation of PigU antisense RNA probe from plasmid |
| Generation of PigU antisense RNA probe from plasmid |
| Generation of PigX antisense RNA probe from plasmid |
| Generation of PigX antisense RNA probe from plasmid |
| Generation of Pgap2 antisense RNA probe from plasmid |

Generation of Pgap2 antisense RNA probe from plasmid
For Sanger Sequencing of Clpex locus in Clpex line
For Sanger Sequencing of Clpex locus in Clpex line
